# Functional connectivity gradients depend on cortical sampling position and brain state

**DOI:** 10.64898/2026.07.14.738484

**Authors:** Muwei Li, Zhaohua Ding, John C Gore

## Abstract

Functional-connectivity analyses often treat signals sampled from nearby cortical positions as interchangeable, despite anatomical and hemodynamic transitions at the gray-white boundary. We tested whether sampling position constitutes a state-sensitive dimension of macroscale functional organization. In 176 Human Connectome Project young adults with four 7 T resting-state and four movie-watching runs, BOLD time series were sampled from subject-specific midthickness and gray-white-boundary surfaces, summarized across 400 cortical parcels, and embedded in a common functional-gradient space. Movie viewing reduced principal-gradient separation between surfaces, but this effect was spatially heterogeneous: Visual, Limbic and Default networks converged, whereas Salience/ventral-attention cortex differentiated. Shapley decomposition and counterfactual analyses showed that the state change reflected coordinated reconfiguration of both surface representations rather than one surface approaching the other. Changes in the original connectivity profiles tracked gradient changes, indicating that the result was not solely introduced by embedding. Both surfaces also exhibited stimulus-locked intersubject functional connectivity during movies. However, stimulus-shared separation captured only part of movie-state organization and did not explain the movie-minus-rest change. Finally, in 172 matched participants, resting-state separation maps generalized from 7 T to 3 T, whereas absolute magnitude and cross-field individual ranking were less stable. These findings establish cortical sampling position as an interpretable measurement dimension of functional-connectivity geometry. Its spatial organization is reproducible across acquisitions, but its magnitude and state dependence vary, emphasizing that where BOLD signals are sampled relative to the cortical boundary can shape conclusions about macroscale brain organization.

## INTRODUCTION

The principal functional-connectivity gradient (G1) is widely treated as a coordinate system of human cortical hierarchy. Yet every such coordinate system begins with a choice of where the BOLD signal is sampled. Its dominant axis extends from primary sensorimotor systems to transmodal association cortex, compactly situating the default-mode network within macroscale organization (Bernhardt et al., 2022; Huntenburg et al., 2018; Margulies et al., 2016). Functional gradients also intersect cortical microstructure, with the two increasingly dissociated toward transmodal cortex (Paquola et al., 2019). Moreover, this organization is state-sensitive: higher-order regions shift their gradient position with task demands, and naturalistic viewing reveals reliable hierarchies that differ from rest (Gao et al., 2022; Kringelbach et al., 2023; Samara et al., 2023). Emerging layer-resolved methods and multilayer analyses further show that resting-state connectivity can differ across cortical depths (Huber et al., 2021; Kotlarz et al., 2025). These studies sharpen, but do not answer, the question addressed here: do adjacent midthickness (core layer of the cortex) and gray-white-boundary samples from conventional whole-brain BOLD acquisitions yield related but nonredundant macroscale gradients, and does their relationship differ between rest and sustained naturalistic viewing?

The gray-white interface is both scientifically consequential and methodologically delicate. It marks a transition between cortex and superficial white matter, where tissue composition, fiber geometry, and vascular contributions change. Adjacent surface samples are therefore neither independent tissue compartments nor necessarily interchangeable: the finite spatial resolution of gradient-echo BOLD, partial-volume mixing, cortical geometry, and depth-dependent vascular effects can produce regionally structured differences in the sampled signal (Huber et al., 2021; Markuerkiaga et al., 2016). At the same time, white-matter BOLD fluctuations exhibit organized functional structure and time-locked responses to stimulation (Gore et al., 2019; Li et al., 2019; Schilling et al., 2023). In our previous study, we quantified functional contrast across this interface using paired gray-and white-matter locations aligned perpendicular to the boundary. Gray-white functional connectivity and the gray-white BOLD power ratio displayed distinct cortical topographies and relationships with myelination, long-range connectivity, neurochemical distributions, and development (Li et al., 2025). Notably, cross-boundary coupling tracked the sensorimotor-association axis, raising the possibility that boundary-proximal functional contrasts intersect the same hierarchy summarized by connectivity gradients. However, those local coupling and power metrics did not test whether each sampling position yields a distinct whole-connectome geometry or whether that geometry varies with brain state. The present comparison is not a laminar measurement; it changes the sampling surface while holding the source volume, participants, and parcellation constant.

Comparing REST and MOVIE provides an informative test of whether this sampling contrast is invariant across acquisition contexts. If cross-surface differences were dominated by static anatomy or a fixed sampling offset, their separation would be expected to remain comparatively stable. A network-specific difference would instead demonstrate non-invariance across acquisitions, although the fixed HCP acquisition order cannot distinguish movie state from session/order, fatigue, or arousal effects and does not identify a neuronal or vascular mechanism. Movies engage visual, auditory, linguistic, social, mnemonic, and executive systems simultaneously, producing distributed functional architectures and reliable changes in connectivity and brain-state dynamics (Demirtas et al., 2019; Rajimehr et al., 2024; van der Meer et al., 2020). For connectivity analysis, the retained multi-minute movie clips reduce the frequency of experimenter-imposed stimulus-to-baseline transitions relative to designs built from repeatedly alternating brief events, and they permit covariance to be estimated during sustained stimulation. Movie-derived connectivity can also carry behaviorally relevant information over short acquisitions (Finn and Bandettini, 2021). This advantage does not eliminate a separate concern: shared evoked responses can systematically inflate within-subject task-connectivity estimates (Cole et al., 2019).

Here, we ask whether macroscale cortical connectivity is preserved across subject-specific midthickness and gray-white-boundary sampling surfaces, and whether their relationship differs between rest and movie viewing. We derive both regional time series from the same preprocessed BOLD volumes, summarize connectivity within the Schaefer 400-region cortical parcellation (Schaefer et al., 2018), and align both representations to a fixed common gradient space (Vos de Wael et al., 2020). We hypothesized that the two surfaces would yield related but non-identical gradients, that their separation would show network-specific REST-MOVIE differences rather than a uniform global offset, and that the resting-state spatial pattern would show cross-acquisition correspondence at 3 T. Complementary analyses algebraically decompose the surface contributions, test whether the gradient pattern is recapitulated in connectivity-profile space, and use intersubject functional connectivity to assess stimulus-shared organization (Nastase et al., 2019; Simony et al., 2016). We also examine time-resolved movie features as a secondary analysis. Throughout, we treat surface differences as measurement-level functional contrasts and do not assign them directly to cortical layers, axonal transmission, or a specific vascular mechanism.

We test these hypotheses using 7 T resting-state and movie fMRI from the Human Connectome Project as the discovery dataset, with same-participant 3 T resting-state data providing a cross-acquisition comparison (Elam et al., 2021). The two sampling surfaces showed a broadly conserved principal-gradient architecture superimposed on network-specific REST-MOVIE differences: average cross-surface separation was lower in the movie acquisition, with convergence in visual, limbic, and default systems but differentiation in salience/ventral-attention cortex. Connectivity-profile analyses recapitulated the principal spatial pattern. Stimulus-shared organization resembled the ordinary movie-acquisition topology without explaining the REST-MOVIE difference. The resting-state spatial pattern showed correspondence across 7 T and 3 T acquisitions despite non-equivalent magnitudes. These findings identify surface-sampling location as a measurable dimension of the aligned G1, whose group topology differed between the REST and MOVIE acquisitions, without assigning the observed contrasts to cortical layers or a specific neural or vascular mechanism.

## METHODS

### Participants, MRI acquisition, and preprocessing

We analyzed publicly available data from the Human Connectome Project Young Adult (HCP-YA) S1200 release. The Washington University Institutional Review Board approved the original data collection, and all participants provided written informed consent. No new human data were acquired for this study. The 7 T discovery sample comprised all 176 participants (70 male and 106 female; age range, 22-35 years) with complete data for four resting-state (REST) and four movie-watching (MOVIE) runs, yielding 704 runs per acquisition context. The same-participant 3 T comparison comprised 172 of these participants with complete, full-length REST1 LR and RL runs. Three participants lacked a required 3 T run and one participant had a shortened LR run (922 rather than 1,200 volumes).

HCP acquisition and reconstruction procedures have been described in detail elsewhere (Elam et al., 2021; Van Essen et al., 2013, 2013; Vu et al., 2017). Briefly, 7 T functional images were acquired with multiband gradient-echo EPI (TR = 1,000 ms, TE = 22.2 ms, flip angle = 45 degrees, multiband factor = 5, 1.6-mm isotropic resolution). Each participant completed four approximately 15-min REST runs (900 volumes each) and four MOVIE runs (901-921 volumes) with AP or PA phase encoding. The eight analyzed runs comprised REST1-PA, REST2-AP, REST3-PA, REST4-AP, MOVIE1-AP, MOVIE2-PA, MOVIE3-PA, and MOVIE4-AP. These runs were acquired across four sessions: Session 1 included REST1 followed by MOVIE1 and MOVIE2; Session 2 included REST2 and retinotopy; Session 3 included REST3 and diffusion imaging; and Session 4 included REST4 followed by MOVIE3 and MOVIE4. The 3 T validation used REST1 LR and RL data acquired with multiband gradient-echo EPI (TR = 720 ms, TE = 33.1 ms, flip angle = 52 degrees, multiband factor = 8, 2-mm isotropic resolution, 1,200 volumes per run). The subject-specific cortical surfaces were derived from the HCP 0.7-mm structural protocol.

We used HCP minimally preprocessed and ICA-FIX-cleaned volumetric functional images. The HCP pipelines correct gradient nonlinearity, head motion and EPI distortion; register functional images to structural images of each participant and standard space; reconstruct cortical surfaces; and remove structured noise with ICA-FIX (Glasser et al., 2013; Salimi-Khorshidi et al., 2014). For the additional preprocessing specific to this study, Connectome Workbench volume-to-surface-mapping was applied separately to midthickness and gray-white-boundary surfaces of each participant. A linear trend and intercept were removed independently from every vertex time series. For the primary REST-MOVIE and 3 T REST analyses, vertex time series were then filtered at 0.01-0.1 Hz, using sampling intervals of 1.0 s at 7 T and 0.72 s at 3 T. Vertex signals were not z-scored before regional averaging. Signals were averaged within the 400 cortical parcels of the Schaefer atlas and summarized by the corresponding seven Yeo networks (Schaefer et al., 2018; Thomas Yeo et al., 2011). Separate detrend-only MOVIE time series were retained for the ISFC and time-resolved feature analyses to preserve stimulus-linked temporal variation. Vertex counts were verified for every parcel and surface, and all retained ROI time series were finite and nonconstant.

### Surface-specific connectivity gradients and the primary REST-MOVIE comparison

The primary analysis compared REST and MOVIE while matching the amount and acquisition direction of data as closely as possible. We retained 13 prespecified movie clips and excluded fixation periods, a short 64-s clip, and repeated Vimeo material. Because the BOLD response follows a stimulus with a delay, the first 5 s after each retained clip began were discarded. REST data were divided using the same clip-duration templates. MOVIE1, MOVIE2, MOVIE3, and MOVIE4 were paired with REST2, REST1, REST3, and REST4, respectively, to maximize phase-encoding agreement.

For each participant, sampling surface, and retained block, we calculated Pearson correlations among the 400 parcel time series. The resulting 400 x 400 functional-connectivity (FC) matrix describes how strongly every pair of parcels fluctuated together. Correlations were converted to Fisher z values before averaging. Within each run, longer blocks received greater weight according to their effective number of observations (n - 3); the four run-level matrices then received equal weight so that no run dominated the participant-level estimate. Mean RelativeRMS displacement for each HCP run was retained as the measure of head motion.

We used functional gradients to summarize each 400 x 400 FC matrix in a small number of continuous spatial axes. Parcels with similar whole-brain connectivity profiles are placed near one another along these axes, whereas parcels with dissimilar profiles are placed farther apart. Gradients were estimated with BrainSpace (Vos de Wael et al., 2020) using a normalized-angle affinity matrix and diffusion-map embedding (90% sparsity, alpha = 0.5, diffusion time = 0, five components, random state = 1). The first gradient (G1) was the primary axis because it captures the dominant sensorimotor-to-association organization described in prior work (Margulies et al., 2016).

Gradient coordinates can be reflected, rotated, or scaled across separate model fits even when their biological organization is similar. We therefore aligned every gradient to one fixed coordinate system before comparing states or surfaces. To construct this reference without favoring a particular state or surface, participant FC matrices were first averaged to obtain four group matrices: REST midthickness, REST white boundary, MOVIE midthickness, and MOVIE white boundary. Their equal-weight grand mean provided an initial reference. The four group gradients were aligned to that reference with Procrustes transformation, which changes orientation but preserves the relative geometry among parcels, and their average formed the locked five-component consensus reference. Every final group and participant gradient was estimated independently and aligned once to this locked reference. We assessed whether alignment was adequate by correlating each aligned component with its corresponding reference component and by calculating the normalized residual difference after alignment. G1 was used for the primary analysis. Distances based on G1-G2 and G1-G3 were examined only as sensitivity analyses because the higher components were less consistently aligned.

For each participant, state, and surface, G1 values were standardized across the 400 parcels. This step removed arbitrary differences in the overall scale of separately estimated gradients while preserving each participant’s spatial pattern. For parcel i, surface separation was defined as Sep_i_ = |G1_midthickness,i_ - G1_white-boundary,i_|. We then calculated ΔSep_i_ = Sep_MOVIE,i_ - Sep_REST,i_. A negative ΔSep therefore means that the two surface representations were closer together during MOVIE (convergence), whereas a positive value means that they were farther apart (differentiation). IntegrationGain was defined as-ΔSep so that positive values indicate convergence. Whole-cortex and network-level scores were obtained by averaging parcel values across all 400 parcels or within each Yeo network.

We first tested whether mean whole-cortex ΔSep differed from zero with a paired t test and summarized its standardized magnitude with Cohen’s d_z_. Because head motion could differ between REST and MOVIE, the primary global and network tests also regressed ΔSep on centered MOVIE-minus-REST RelativeRMS of each participant. In this model, the intercept estimates the mean state-related change after accounting for motion. Seven network tests were performed, so their p values were adjusted with the Benjamini-Hochberg false-discovery-rate procedure (Benjamini and Hochberg, 1995). The 95% confidence intervals shown for group estimates were obtained by resampling participants with replacement 5,000 times; they describe uncertainty across participants rather than variability across parcels or runs.

Finally, we asked whether the REST-to-MOVIE change was the same in every network or was spatially heterogeneous. A repeated-measures linear mixed model included condition, network, their interaction, and centered motion as fixed effects. The condition-by-network interaction was the term of interest: a significant interaction indicates that the condition difference varies among the seven networks instead of reflecting a uniform cortex-wide shift. Random intercepts for participant and participant-by-network accounted for the repeated observations contributed by the same individual and for stable differences among that individual’s network-level values.

Several analyses tested whether the primary result depended on a particular scaling choice, motion threshold, or subset of participants. To determine how much of the effect reflected gradient scale rather than spatial topology, we repeated the global comparison with unstandardized aligned G1 distances. A low-motion analysis retained the 145 participants whose mean RelativeRMS across the two conditions was below 0.15. Spatial reproducibility was evaluated with 100 repeated split-half analyses. Each half of the cohort independently rebuilt its group FC matrices and consensus reference, and the two resulting ΔSep maps were correlated; the full-cohort reference was used only to identify the corresponding gradient component.

We also repeated the analysis separately for the four phase-encoding-matched MOVIE-REST run pairs. These estimates show whether the direction of an effect is consistent across runs, but they are repeated measurements from the same participants and were not treated as independent replications. Reliability of the participant-level IntegrationGain score was summarized with intraclass correlation coefficients (ICCs) (McGraw and Wong, 1996). Single-pair ICCs estimate how reproducible the ranking of an individual would be from one matched run pair, whereas four-pair average ICCs estimate reliability after combining all four pairs. We report both absolute-agreement ICCs, which require scores to be numerically similar, and consistency ICCs, which require only preservation of relative ranking.

Because large veins can influence gradient-echo BOLD signals near the cortical boundary, we tested whether a simple group-average venous map could explain the spatial pattern of ΔSep. The VENAT partial-volume atlas was sampled on HCP Q1-Q6_R440 group-average midthickness and white surfaces using the same trilinear Workbench procedure (Huck et al., 2019). For each parcel, venous depth contrast was defined as mean VENAT partial volume at midthickness minus that at the white boundary.

We correlated the venous depth-contrast map with the group ΔSep map using two-sided Spearman correlation; Pearson correlation was secondary. Conventional parcelwise significance tests are inappropriate for cortical maps because neighboring parcels are spatially similar. We therefore compared the observed correlation with 10,000 null correlations generated by rotating the parcel map on the cortical surface while preserving hemispheres and spatial smoothness. Rotated parcel assignments were obtained from spherical parcel centroids using Hungarian matching (Alexander-Bloch et al., 2018). This analysis asks whether the visible-vein topology is sufficient to explain the group-average spatial pattern; it cannot exclude participant-specific anatomy or state-dependent vascular effects.

### How the two sampled surfaces contributed to the state-related change

ΔSep describes the net change in distance between the two surface representations, but it does not reveal how each representation contributed. For example, the same convergence could arise because only the midthickness representation changed, only the white-boundary representation changed, or both changed in a coordinated manner. We addressed this ambiguity with a two-feature Shapley decomposition (Shapley, 1953), a mathematical procedure that assigns the total change to the two surfaces while averaging over the order in which their state changes are considered.

Before calculating distances, each aligned component was standardized across parcels within participant, state, and surface. For analyses using k = 1, 2, or 3 components, distance was the parcelwise Euclidean distance divided by √k, which kept values comparable as components were added. Four distances represented the observed and counterfactual configurations listed in the table below: D_00_ compared both REST representations; D_11_ compared both MOVIE representations; D_10_ changed only midthickness to its MOVIE configuration; and D_01_ changed only the white boundary to its MOVIE configuration.

| Distance | Midthickness | White<br>Boundary | Meaning |
| --- | --- | --- | --- |
| **D<sub>00</sub>** | REST | REST | Observed REST configuration |
| **D<sub>10</sub>** | MOVIE | REST | Counterfactual configuration in which only the midthickness surface is changed to the MOVIE state |
| **D<sub>01</sub>** | REST | MOVIE | Counterfactual configuration in which only the white-boundary surface is changed to the MOVIE state |
| **D<sub>11</sub>** | MOVIE | MOVIE | Observed MOVIE configuration |

D_00_ and D_11_ are the two observed configurations, whereas D_10_ and D_01_ are mixed-state counterfactuals used only for attribution. The midthickness contribution was Φ_mid_ = 0.5[(D_10_ - D_00_) + (D_11_ - D_01_)], and the white-boundary contribution was Φ_white_ = 0.5[(D_01_ - D_00_) + (D_11_ - D_10_)]. These two terms sum exactly to D_11_ - D_00_, which is ΔSep. A negative contribution means that the corresponding surface reduced cross-surface distance and promoted convergence; a positive contribution means that it increased distance and promoted differentiation. The terms are algebraic attributions in gradient space and do not imply anatomical movement, causal direction, or one surface driving the other. For a more direct illustration, we also calculated D_10_ - D_00_ and D_01_ - D_00_. These contrasts ask what would happen to separation if one surface adopted its MOVIE configuration while the other remained fixed in its REST configuration. A negative value means that changing that surface alone would bring the representations closer together; a positive value means that it would move them farther apart. Participant-level network contributions were tested against zero. State-distance contrasts were additionally adjusted for the REST-to-MOVIE motion difference, and BH-FDR was applied separately within each seven-network family. The G1 decomposition was primary. G1-G2 and G1-G3 distances, estimates for all networks, the direct counterfactual contrasts, and group network trajectories in G1-G2 space were used to test dimensional robustness or aid visualization.

We next asked whether the spatial pattern of functional separation followed a broad cortical microstructural gradient. Structural context was represented by the HCP bias-corrected myelin map, an unsmoothed T1w/T2w-derived contrast that is sensitive to myelin but is not a direct measure of myelin content (Glasser et al., 2013; Glasser and van Essen, 2011). The map of each participant was averaged within Schaefer parcels and standardized across the 400 parcels; the primary group map was the mean of these standardized maps.

Two-sided Spearman spin tests related the group structural map to four prespecified functional maps: REST separation, ΔSep, Φ_mid_, and Φ_white_. The same 10,000 bilateral rotations controlled for cortical spatial autocorrelation, and BH-FDR was applied across the four outcomes. Analyses using the unstandardized mean map tested sensitivity to scaling, and a paired-spin test compared the structural associations of Φ_mid_ and Φ_white_. Additional sensitivity analyses asked whether a quadratic relationship improved on a linear one, whether slopes differed among networks beyond their mean levels, and whether within-participant associations exceeded those obtained with a leave-one-participant-out group map. These analyses provided structural context but were not used to claim an individual-specific myelin mechanism.

### Connectivity-profile analysis independent of gradient embedding

Gradient embedding is a nonlinear summary of the original FC matrix. We therefore tested whether the same REST-to-MOVIE pattern was visible directly in the connectivity profiles from which the gradients were derived. For each source parcel, its FC profile was the row of correlations linking it to the other 399 parcels. Midthickness and white-boundary profiles were standardized separately across targets, and the self-connection was excluded.

For each source parcel, cross-surface profile distance was the root-mean-square difference between its two standardized 399-target profiles. Parcel distances were averaged within each source network to obtain seven whole-profile summaries. To identify which target systems contributed, we also divided every source profile into the seven target networks, producing 49 source-by-target combinations. For both summaries, the state contrast was MOVIE minus REST, so negative values indicate profile convergence and positive values indicate profile differentiation. Whole-profile and source-by-target contrasts were tested against zero both before and after adjustment for the REST-to-MOVIE motion difference. BH-FDR was applied across the seven source-network tests or across all 49 source-by-target tests, respectively. The 95% confidence intervals plotted for group means were generated by 5,000 participant bootstrap resamples.

We then asked whether participants who showed stronger FC-profile convergence also showed stronger G1 convergence. Within each source network, FC-profile change and G1 ΔSep were each adjusted for motion, and the residual values were correlated across participants. BH-FDR was applied across the seven correlations. Because this analysis uses the original FC rows rather than the embedded gradients, agreement between the measures shows that the gradient result has a corresponding pattern in the underlying connectivity geometry; it does not establish that one measure causes the other.

### Stimulus-shared connectivity and time-resolved movie features

Conventional FC is calculated within each participant and therefore combines movie-locked responses with spontaneous or idiosyncratic fluctuations. To isolate the portion of connectivity that was temporally shared across viewers, we calculated intersubject functional connectivity (ISFC) from the detrend-only MOVIE time series (Nastase et al., 2019; Simony et al., 2016). For each participant, ISFC correlates a parcel’s time series with the corresponding signal averaged across all other participants. Fluctuations that occur at different times in different viewers tend to cancel, whereas responses consistently locked to the movie are retained. ISFC is therefore an index of stimulus-shared organization, not a pure measure of direct neural interaction.

Only the linear trend was removed for ISFC; band-pass filtering was omitted to avoid attenuating movie-related fluctuations outside the conventional resting-state band. We used the same 13 clips and windows from 5 s after onset to clip offset. Within participant, clip, parcel, and surface, time series were standardized across TRs. Group ISFC was calculated as the exact mean of ordered cross-participant correlations. Clip matrices were converted to Fisher z values, weighted by effective duration within each run, and then combined so that each of the four runs contributed one quarter of the final estimate. The diagonal was set to zero before gradient embedding so that same-parcel correlations across viewers could not dominate the similarity among whole-brain profiles.

We first tested whether ISFC was stronger than expected without shared movie timing. Parcel signals were averaged within the seven Yeo networks, and the summary statistic was the mean absolute ISFC among different networks. For each of 1,000 null repetitions, the time series of every participant was circularly shifted within each clip by at least 20 s, using the same shift for both surfaces. Circular shifting preserves each signal autocorrelation and amplitude distribution while breaking its alignment with other viewers and with the movie. One-sided empirical p values tested whether observed ISFC exceeded this null distribution.

We then embedded the final run-balanced parcelwise ISFC matrices with the same BrainSpace settings and aligned the gradients to the locked 7 T reference. After standardizing G1 across parcels, we calculated midthickness-to-white-boundary separation. Spin tests assessed whether the resulting ISFC separation map resembled ordinary within-participant MOVIE separation or the primary ΔSep map. All reported ISFC inferences use this run-balanced analysis. The earlier equal-clip map is shown only as a weighting-scheme sensitivity analysis, whereas an earlier post-embedding circular-shift null is shown only to document why that null was discarded; neither alternative supplies the reported inference.

A separate time-resolved analysis asked whether the two surfaces became more similar in their stimulus-locked responses at moments of semantic change. For every participant, parcel, and clip, standardized midthickness and white-boundary signals were transformed into a common mode, (mid + white)/√2, and a differential mode, (mid - white)/√2. The common mode captures fluctuations expressed similarly at both surfaces, whereas the differential mode captures fluctuations that distinguish them. We calculated instantaneous synchrony across participants for both modes and defined depth commonality as common-mode synchrony minus differential-mode synchrony. Positive values therefore indicate that shared cross-viewer activity was expressed more similarly across the two surfaces. Parcel values were averaged for the whole cortex and for each Yeo network.

Semantic features were obtained from the HCP WordNet resource (Huth et al., 2012; Miller, 1995). Semantic change at each TR was defined as one minus the cosine similarity between consecutive WordNet vectors, so larger values indicate a larger shift in the represented semantic content. Semantic density was the proportion of nonzero WordNet entries. Low-level visual controls were the root-mean-square MotionEnergy level and its root-mean-square temporal change (Nishimoto et al., 2011). Feature time series were shifted 5 s earlier than the BOLD measurements to approximate the hemodynamic delay.

Predictors and depth-commonality outcomes were standardized within each clip so that the model estimated moment-to-moment associations rather than differences in average level among clips. For the whole cortex and each of the seven networks, we estimated the association between semantic change and depth commonality after removing variation related to semantic density, MotionEnergy level, MotionEnergy change, and clip identity. The semantic coefficient therefore asks whether moments containing larger semantic transitions showed greater cross-surface commonality than other moments in the same clips with comparable low-level visual and semantic-density features. Incremental R^2^ quantified the additional variance explained by semantic change beyond those controls.

Statistical significance was evaluated with 1,000 independent circular shifts of semantic change within each clip, each by at least 20 s. This null preserves clip membership and temporal autocorrelation but breaks the moment-to-moment alignment between semantic change and the BOLD outcome. Two-sided empirical p values were corrected across the whole-cortex and seven network outcomes. To assess whether pooled effects generalized across movie content, controlled coefficients were also estimated separately for the 13 clips and tested across clips. Leave-one-clip-out models, predictor correlations, and lags from 0 to 10 s were supporting sensitivity analyses rather than independent confirmatory tests.

### Cross-acquisition validation of REST organization at 3 T

The 3 T analysis asked whether the spatial organization observed during 7 T REST could be recovered under a different acquisition protocol in the same participants. It did not test the MOVIE effect and was not an independent-cohort replication. For each of 172 participants, 3 T REST1 LR and RL FC matrices were calculated separately from the band-pass-filtered parcel time series and combined by an equal-weight Fisher-z mean. We rebuilt these matrices from the parcel time series rather than reusing earlier 3 T derivatives.

Fresh 3 T session matrices and the matched four-run 7 T REST matrices were embedded independently. All gradients were aligned to the reference locked during the 7 T discovery analysis; the 3 T data did not help define this primary coordinate system. This design tests whether 3 T organization corresponds to an already established 7 T pattern rather than allowing both datasets to determine a new shared solution. Components were standardized across parcels, and separation across the first k components was the Euclidean midthickness-to-white-boundary distance divided by √k. G1 was primary, with G1-G2 and G1-G3 used as dimensional sensitivity analyses.

Spatial generalization was evaluated by averaging G1 separation of each participant at every parcel and correlating the resulting 3 T and 7 T group maps. Significance was assessed with 10,000 bilateral spin permutations, which preserve the broad spatial smoothness of cortical maps (Alexander-Bloch et al., 2018). Spearman correlation and Pearson correlation after removing the seven-network means of each map tested whether correspondence was robust to rank-based estimation and extended beyond broad differences among networks.

Additional sensitivity analyses used gradients estimated from group-average FC matrices and a symmetric coordinate system constructed with equal contributions from matched 3 T and 7 T group embeddings. The symmetric analysis was not the primary test because it allows the validation dataset to influence the coordinate system; the established 7 T reference was used only to identify component orientation. For every participant and surface, we also inspected reference correlations and alignment residuals to identify poorly aligned estimates.

Group-map similarity does not guarantee that participants retain the same relative ranking across field strengths. We therefore correlated whole-cortex 3 T and 7 T separation of each participant before and after adjusting each measure for its own mean RelativeRMS. Significance was assessed with 10,000 participant-label permutations, which ask whether the observed pairing of 3 T and 7 T values is stronger than random pairing. The 95% confidence interval was obtained from 5,000 participant bootstrap samples. Family relatedness was not modeled, so permutations were unrestricted rather than performed within family blocks.

We separately asked whether the absolute magnitude of separation differed between 3 T and 7 T. The within-participant difference was tested with a paired t test and summarized with Cohen’s d_z_. A complementary regression estimated the motion-adjusted mean difference using HC3 robust standard errors, which reduce sensitivity to unequal residual variance (MacKinnon and White, 1985). Seven-network means described whether broad spatial organization was preserved despite any global magnitude shift. Finally, LR-RL correlations and two-way absolute-agreement ICCs quantified the reliability of a single 3 T phase-encoding run [ICC(A,1)] and of the average of LR and RL [ICC(A,2)] (McGraw and Wong, 1996). Together, these analyses distinguish spatial-map correspondence, participant ranking, numerical equivalence, and within-3 T reliability rather than treating them as the same form of validation.

## RESULTS

### Movie viewing reconfigures cross-surface functional-gradient organization

We first defined a complete-case 7 T discovery sample in which every participant contributed all four REST and all four MOVIE runs. Of 184 candidates, 176 met this requirement, providing 704 runs in each acquisition context (Fig. S1a). Thirteen long-form movie clips were retained after excluding fixation periods, a 64-s source clip, and repeated Vimeo material (Fig. S1b). For each participant, BOLD signals sampled at the midthickness and white boundary were summarized in 400 cortical parcels and converted to duration-matched FC matrices (Fig. S1c). The prespecified MOVIE-REST run pairings produced closely matched effective durations after the 5-s onset buffer (Fig. S1d). A subset of 172 participants also had complete 3 T REST data and was reserved for the cross-acquisition analysis (Fig. S1e).

The primary question was whether the distance between the two sampled-surface representations changed between REST and MOVIE within the common gradient coordinate system (Fig. 1a). All 400 parcels had adequate vertex coverage (Fig. S2a). The participant-level G1 gradients corresponded closely to the locked reference for both conditions and surfaces, indicating that the same dominant connectivity axis was being compared (Fig. S2b). Correspondence was strongest for G1-G2, remained acceptable for G3, and weakened for G4-G5 (Fig. S2c); residual differences after group alignment were modest (Fig. S2d). We therefore treated standardized G1 separation as the primary measure and used higher-dimensional distances only as sensitivity analyses.

**Figure 1.**
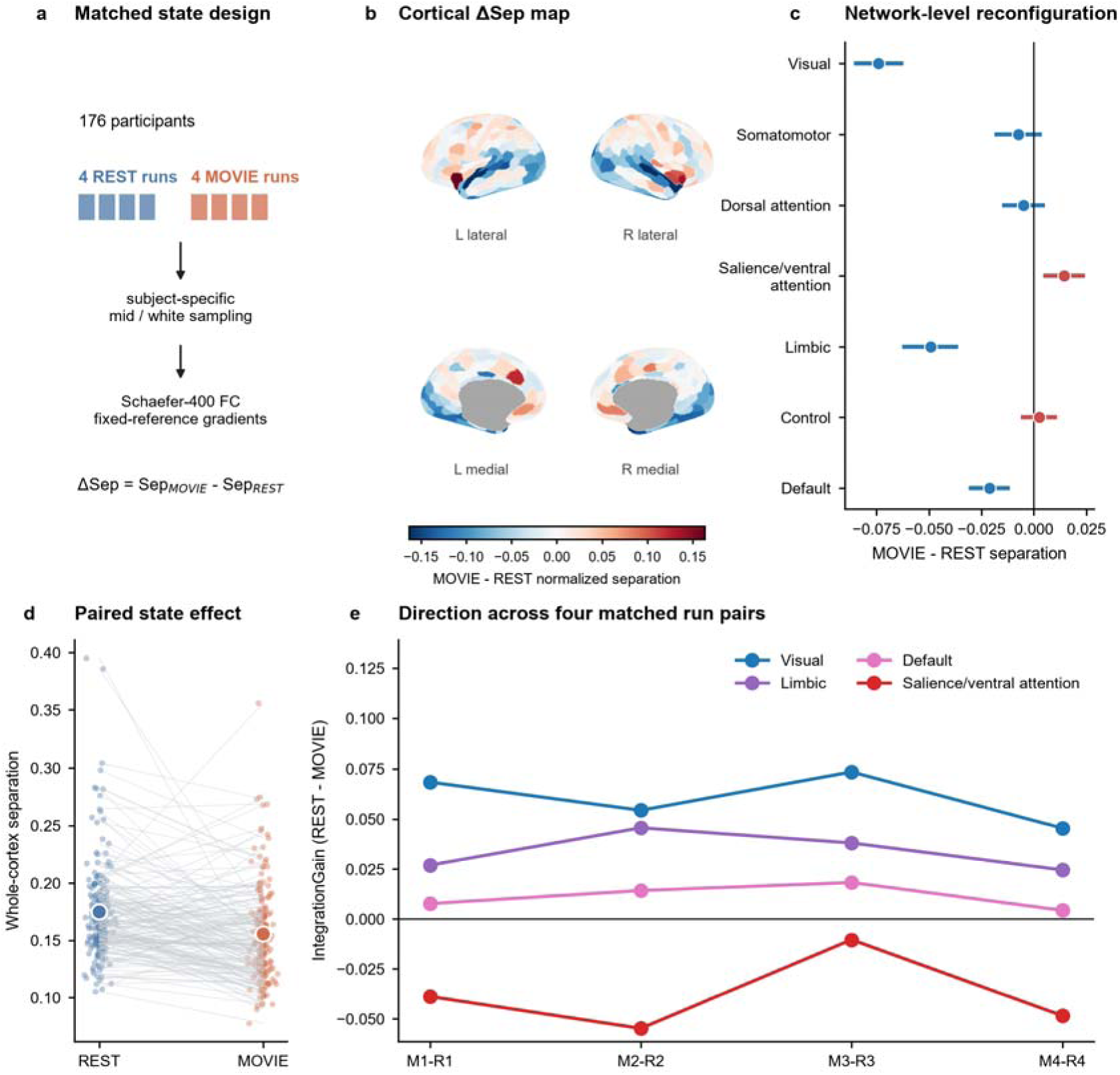
Movie viewing induces network-specific cortical depth-gradient reconfiguration. a, Matched-state analysis of 176 participants with four REST and four MOVIE runs. BOLD signals were sampled from subject-specific midthickness and gray-white-boundary surfaces, reduced to Schaefer-400 cortical parcels, converted to duration-matched functional-connectivity matrices and embedded using a fixed common gradient reference. Separation, Sep, is the within-subject scale-normalized absolute G1 distance between the two sampled surfaces. ΔSep is Sep(MOVIE) minus Sep(REST); negative values indicate cross-depth convergence and positive values differentiation. b, Across-participant mean parcelwise ΔSep displayed on fsLR-32k very-inflated surfaces. Blue denotes convergence, red differentiation and gray the medial wall without a Schaefer assignment. c, Seven-network mean ΔSep. Points are group means and horizontal intervals are nonparametric bootstrap 95% confidence intervals; the vertical line denotes no state change. Blue and red encode effect direction, not statistical significance. Network inference used a linear model adjusting for the MOVIE-minus-REST motion difference and BH-FDR across seven networks. Visual, Limbic and Default showed significant convergence, whereas Salience/ventral attention showed significant differentiation. d, Paired whole-cortex separation for all 176 participants. Translucent points are participants, gray lines join the same participant and large circles are group means. Mean separation decreased from 0.175 in REST to 0.155 in MOVIE (paired d_z_=-0.40; motion-adjusted p=3.21 × 10^-7^). e, IntegrationGain across four prespecified duration-matched run pairs for the four networks surviving the panel-c motion-adjusted FDR analysis. IntegrationGain is Sep(REST) minus Sep(MOVIE), the negative of ΔSep; positive values therefore indicate convergence and negative values differentiation. Run-pair estimates assess directional consistency within the same participants and are descriptive, not four independent replications.

The REST-to-MOVIE change was not uniform across cortex. The parcelwise ΔSep map showed broad convergence in visual, limbic, and default territories, together with focal differentiation in salience/ventral-attention cortex (Fig. 1b). After accounting for the motion difference and correcting across seven networks, ΔSep was negative in Visual (-0.074, q = 2.56 × 10^-27^), Limbic (-0.049, q = 7.57 × 10^-13^), and Default (-0.021, q = 8.56 × 10^-6^), but positive in Salience/ventral attention (0.015, q = 0.0022); the other three networks were not significant (Fig. 1c). The condition-by-network interaction directly tested whether these condition differences varied among networks and was strongly significant (F(6,2449) = 39.47, p = 4.85 × 10^-46^; Fig. S3c). Thus, the global tendency toward convergence comprised distinct and sometimes opposing network responses rather than uniform compression.

Averaged across all 400 parcels, normalized separation decreased from 0.175 during REST to 0.155 during MOVIE (ΔSep =-0.020; paired d_z_ =-0.40; motion-adjusted p = 3.21 × 10^-7^; Fig. 1d). The effect was larger before standardizing G1 scale (d_z_ =-0.74), showing that changes in gradient scale contributed to the raw difference but did not account for the normalized convergence (Fig. S3a). Although ΔSep was weakly related to the REST-to-MOVIE motion difference (r = 0.17), all primary participant-level REST-to-MOVIE models adjusted for this motion difference, and the low-motion subset retained a similar negative mean (n = 145, mean = −0.018; Fig. S3b). Across 100 independently rebuilt split halves, the parcelwise ΔSep map was reproducible (median map correlation approximately 0.72; Fig. S3d).

The network directions were also broadly consistent across the four matched run pairs. Visual, Limbic, and Default showed convergence in all four pairs, whereas Salience/ventral attention showed differentiation in all four (Fig. 1e). The whole-cortex effect itself was pronounced only in the third pair, emphasizing that run pairs provide repeated checks of direction within the same participants rather than four independent replications (Fig. S3e). Participant rankings were poorly reproducible from a single run pair and improved when all four pairs were averaged, but even the four-pair ICCs were too modest to support interpretation as a stable individual trait (Fig. S3f).

We then examined whether the group-average pattern simply followed where visible veins are more prominent at one sampling surface than the other. The VENAT atlas produced a structured midthickness-minus-white venous map (Fig. S8a), but that map was spatially distinct from ΔSep (Fig. S8b). Their association was weak (Spearman ρ =-0.080; Fig. S8c) and was not unusual relative to 10,000 spatial rotations (p_spin_ = 0.545; Fig. S8d). The visible-vein group topology was therefore insufficient to explain the observed pattern, although this analysis cannot exclude participant-specific anatomy or state-dependent vascular contributions.

### Both sampled-surface representations contributed to the state-related change

We next asked how the midthickness and white-boundary representations contributed to ΔSep. The four observed and counterfactual distances isolate what happens when the state of one surface is changed while the other is held fixed, and the two Shapley terms allocate the full observed change between the surfaces (Fig. 2a). This is a mathematical decomposition of movement in aligned gradient space; it does not identify anatomical or causal direction.

**Figure 2.**
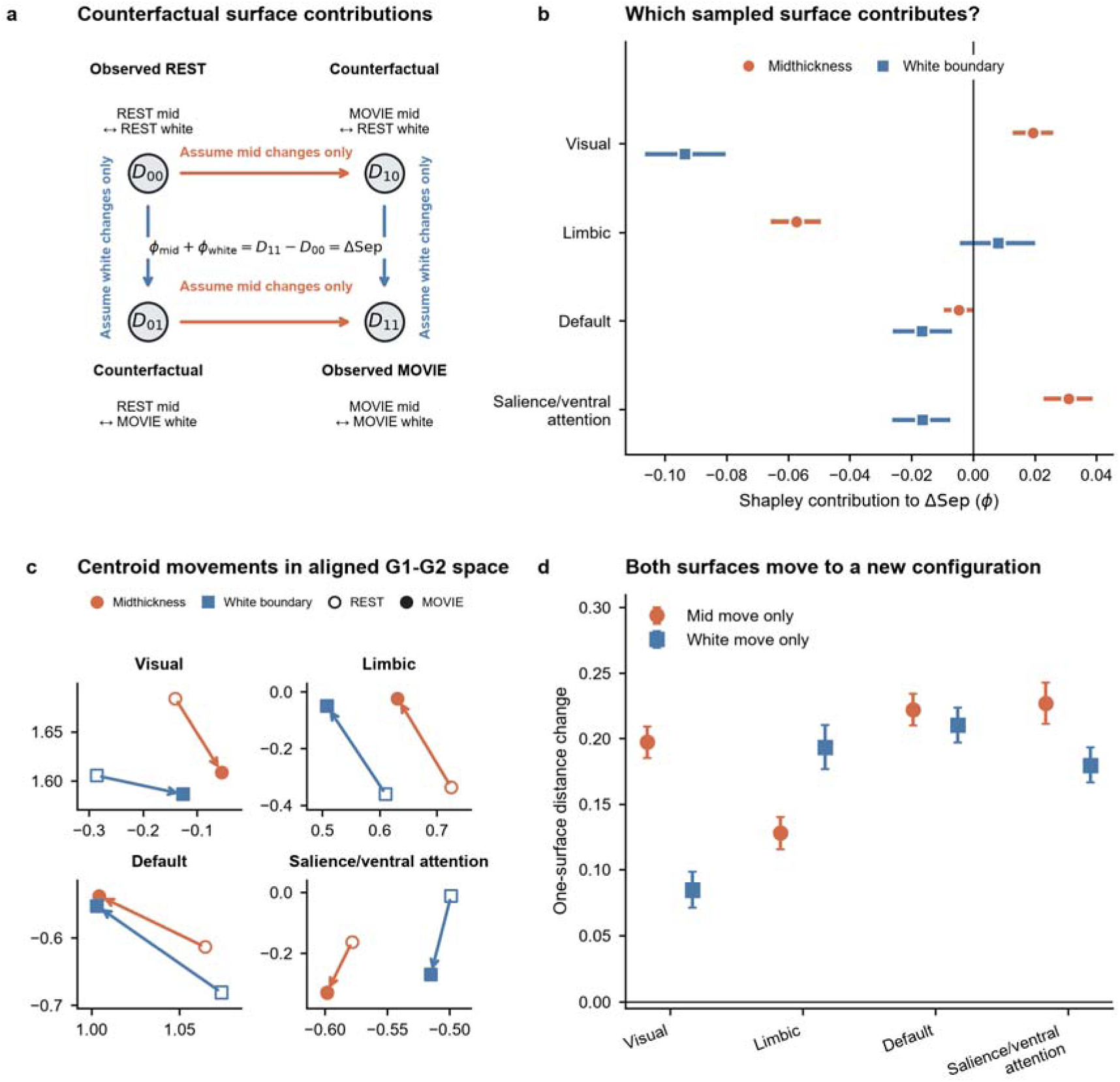
Distinct surface contributions generate convergence and G1-specific differentiation. a, Counterfactual state space used to decompose the observed REST-to-MOVIE separation change. Each D denotes the normalized G1 gradient-space distance between the two surface representations joined by the double arrow: D_00_ compares REST midthickness with REST white boundary, D_10_ compares MOVIE midthickness with REST white boundary, D_01_ compares REST midthickness with MOVIE white boundary and D_11_ compares MOVIE midthickness with MOVIE white boundary. Thus, the double arrow denotes a distance comparison, not addition. Horizontal arrows change midthickness while holding the white boundary fixed; vertical arrows change the white boundary while holding midthickness fixed. D_00_ and D_11_ are observed REST and MOVIE distances, whereas D_10_ and D_01_ are counterfactual mixed-state distances. The two Shapley contributions sum exactly to ΔSep. b, Midthickness Shapley contributions, φ_mid_ (orange circles), and white-boundary Shapley contributions, φ_white_ (blue squares), for the four focal networks; intervals are bootstrap 95% confidence intervals. Negative φ reduces G1 separation and therefore promotes convergence, whereas positive φ increases G1 separation and promotes differentiation; φ_mid_plus φ_white_ equals ΔSep. c, REST-to-MOVIE movements in aligned G1-G2 space; hollow symbols denote REST, filled symbols MOVIE, orange circles midthickness and blue squares the white boundary. Panel b quantifies contributions to G1 separation, whereas panel c visualizes centroid movement along the first and second functional-connectivity gradients; an arrow’s direction along either axis does not directly indicate the sign of φ. d, Direct one-surface-at-a-time distance changes. Positive values mean that changing either surface alone increases its distance from the other surface’s REST representation. Thus, final convergence reflects coordinated repositioning of both representations, not one approaching the other’s original position. These effects are defined in aligned functional-gradient space and do not imply anatomical movement or causal influence.

The contributions differed markedly among networks (Fig. 2b). Visual convergence was dominated by the white-boundary contribution (Φwhite =-0.093, q = 2.16 × 10^-31^), while the positive midthickness contribution partly opposed convergence (Φmid = 0.019, q = 2.57 × 10^-9^). Limbic convergence showed the opposite allocation: it was dominated by midthickness (Φmid =-0.057, q = 4.75 × 10^-33^), with a small nonsignificant white-boundary contribution. Both surfaces contributed to Default convergence, although the white-boundary term was larger (Φmid = - 0.0046, q = 0.035; Φwhite =-0.0166, q = 0.00054). Salience/ventral-attention differentiation was driven by a positive midthickness contribution (Φmid = 0.031, q = 3.09 × 10^-14^) that exceeded an opposing white-boundary contribution (Φwhite =-0.016, q = 0.00054). Dorsal-attention and Control also contained opposing surface contributions that largely cancelled in their net ΔSep values (Fig. S4a). These results show that the same net change in cross-surface separation can arise from distinct, and sometimes opposing, changes in the two surface representations across networks, arguing against a single surface or uniform cortical process as the source of the REST-to-MOVIE effect.

Network centroids in G1-G2 space provided an intuitive view of these attributions: both surface representations changed from REST to MOVIE, but their directions and magnitudes differed by network (Fig. 2c; Fig. S4c). Changing either surface alone (mid move only D_10_ − D_00_, or white move only D_01_ − D_00_) actually increased its distance from the REST representation of the other surface in each focal network (Fig. 2d), a pattern also present across all seven networks (Fig. S4b). Final convergence therefore did not occur because one representation simply approached the other’s original REST position. It emerged only after both representations occupied a new MOVIE configuration.

The dimensional sensitivity analysis clarified the scope of this interpretation. Visual convergence persisted when distance included G1-G2 or G1-G3. In contrast, the positive Salience/ventral-attention effect reversed after additional components were included (Fig. S4d). Salience differentiation is therefore specific to the dominant G1 axis and should not be generalized to the entire low-dimensional connectivity manifold.

The T1w/T2w-derived structural map showed the expected broad cortical organization (Fig. S9a) but did not provide a sufficient spatial explanation for the functional effects. Its association with REST separation was nonsignificant after spin testing (ρ = 0.14, p_spin_ = 0.396, q = 0.396; Fig. S9b), as were its associations with ΔSep (ρ =-0.22, p_spin_ = 0.273, q = 0.364; Fig. S9c), Φmid (ρ = 0.22, p_spin_ = 0.095, q = 0.213; Fig. S9d), and Φwhite (ρ =-0.26, p_spin_ = 0.107, q = 0.213; Fig. S9e). The difference between the two contribution associations was suggestive but did not reach significance (paired-spin p = 0.060). Thus, the group myelin-sensitive contrast did not account for either baseline separation or its state-related change.

### Connectivity-profile changes reproduced the gradient pattern

Because gradients are nonlinear summaries of FC, we asked whether the same pattern was present in the original connectivity profiles. For every source parcel, we compared its standardized connections to the other 399 parcels between the two sampling surfaces (Fig. 3a). Negative MOVIE-minus-REST values mean that the midthickness and white-boundary profiles became more similar during MOVIE; positive values mean that they became more distinct.

**Figure 3.**
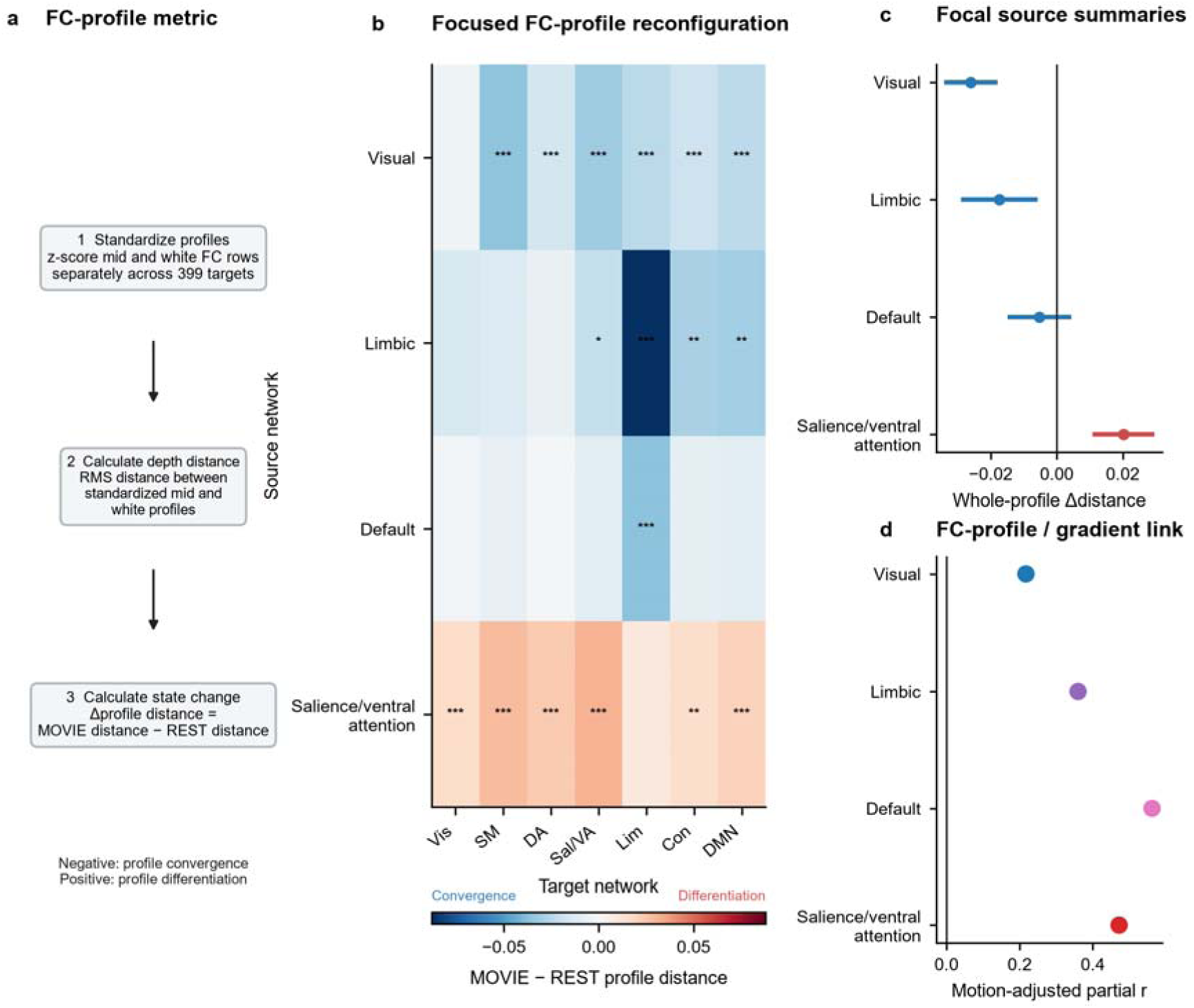
FC-profile reconfiguration provides an embedding-independent basis for the gradient pattern. a, FC-profile distance metric. For every source parcel and condition, midthickness and white-boundary FC rows were separately z-scored across the other 399 cortical targets. Their root-mean-square distance quantified cross-depth profile separation, and the state effect was defined as MOVIE distance minus REST distance. Negative values indicate that the two depth-specific FC profiles become more similar during MOVIE (profile convergence); positive values indicate greater dissimilarity (profile differentiation). b, Target-resolved MOVIE-minus-REST profile-distance changes for the four focal source networks and all seven target networks. Blue denotes convergence and red differentiation. Asterisks denote motion-adjusted BH-FDR across the complete 49 source-target tests. c, Whole-profile source-network summaries with bootstrap 95% confidence intervals. Unlike panel b, these values were calculated directly from each source parcel’s complete 399-target FC profile and were not obtained by averaging the seven network-level heat-map cells. d, Associations between subject-level whole-profile change and G1 separation change after residualizing both variables for the MOVIE-minus-REST motion difference. Positive partial correlations indicate that stronger profile convergence accompanies stronger G1 convergence, and stronger profile differentiation accompanies stronger G1 differentiation. FC-profile correspondence does not establish causality.

The target-resolved analysis reproduced the main network topology (Fig. 3b). Visual sources converged with six of seven target networks after motion-adjusted FDR correction, whereas Salience/ventral-attention sources differentiated with six of seven targets. Limbic sources showed their strongest convergence toward Limbic targets, and Default sources converged selectively toward Limbic targets. Across all 49 source-target combinations, convergence also appeared for Dorsal-attention and Control sources; Salience/ventral attention was the only source network with broadly positive effects (Fig. S5a).

When complete 399-target profile regarding each source parcel was summarized as a single distance, Visual (-0.026, q = 4.30 × 10^-10^), Dorsal attention (-0.021, q = 1.32 × 10^-7^), Limbic (- 0.017, q = 0.0018), and Control (-0.020, q = 1.34 × 10^-6^) converged, whereas Salience/ventral attention differentiated (0.020, q = 1.05 × 10^-5^; Fig. 3c; Fig. S5b). Somatomotor and Default did not show significant whole-profile changes. The null Default average nevertheless contained a selective Default-to-Limbic convergence effect (-0.036, q = 1.04 × 10^-4^; Fig. S5c), showing why a network’s overall profile can remain stable even when particular target relationships change.

Participants with larger FC-profile changes also tended to show larger G1 changes after both measures were adjusted for motion. The partial correlations were positive in Visual (r_partial_ = 0.217), Limbic (0.360), Default (0.562), and Salience/ventral attention (0.472), with all q <= 0.0045 (Fig. 3d). All seven source networks showed positive associations in the complete analysis (range, 0.183-0.584; all survived FDR; Fig. S5d). Thus, the cross-surface gradient pattern had a corresponding change in the underlying connectivity profiles. This agreement supports a shared organizational basis but does not show that profile change causally produced the gradient change.

### Stimulus-shared connectivity and semantic transitions during movie viewing

We next asked which part of MOVIE organization was temporally shared across viewers. Before gradient embedding, observed mean absolute ISFC among Yeo networks was far above the circular-shift null at both surfaces: 0.0361 versus 0.00106 at midthickness and 0.0308 versus 0.00094 at the white boundary (both p = 0.001; Fig. 4a). Thus, both sampling surfaces contained robust connectivity synchronized to the movie. The run-balanced weighting gave longer clips more influence within a run but gave each of the four runs equal total influence (Fig. S6a). The run-balanced map remained highly similar to the alternative equal-clip map (r = 0.979, p_spin_ = 0.00030; Fig. S6b), showing that the spatial topology was robust to this weighting choice. Both run-balanced ISFC G1 gradients also aligned well to the locked reference (r = 0.880 and 0.868; Fig. S6c).

**Figure 4.**
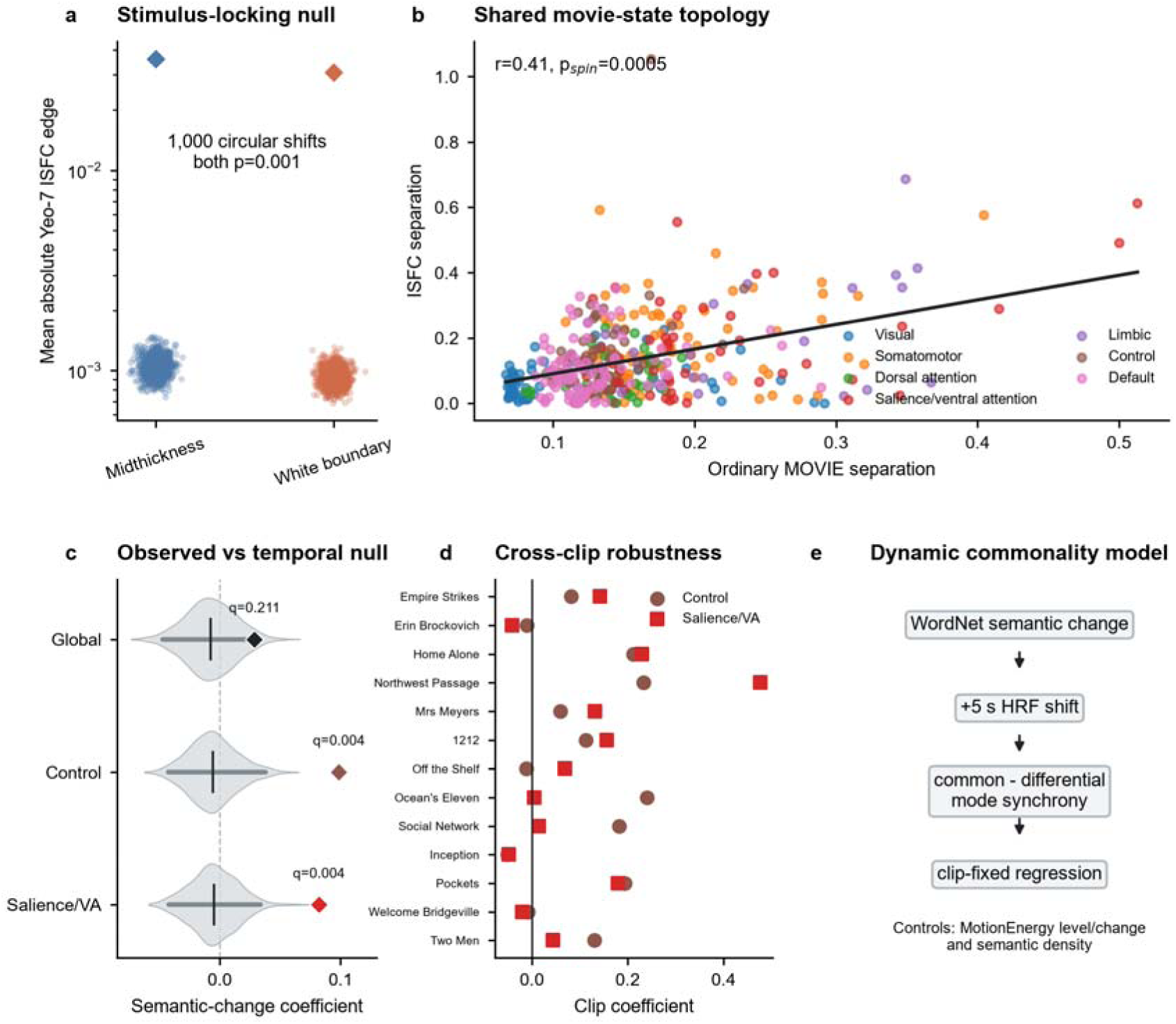
Shared movie timing and semantic transitions organize cross-depth commonality. a, Stimulus-locking test performed on mean absolute Yeo-7 intersubject functional-connectivity (ISFC) edges before gradient embedding. Diamonds are observed values and translucent points are 1,000 null values generated by independently circularly shifting each participant within each clip. Observed midthickness and white-boundary values were 0.0361 and 0.0308, compared with null means of 0.00106 and 0.00094; both one-sided p=0.001. b, Spatial correspondence across 400 cortical parcels. The x axis is normalized G1 midthickness-to-white-boundary separation from ordinary within-participant MOVIE FC; the y axis is the corresponding separation from stimulus-shared ISFC. Colors identify Yeo-7 networks, and the black line is the least-squares fit. Inference used 10,000 hemisphere-preserving spins (Pearson r=0.410, p_spin_=0.00050), supporting a shared movie-state topology rather than showing that ISFC explains MOVIE-minus-REST ΔSep. c, Semantic-change coefficients for the global cortex, Control and Salience/ventral-attention networks. Positive coefficients indicate that larger WordNet semantic transitions, shifted by 5 s, predict greater common-mode relative to differential-mode synchrony after MotionEnergy level/change, semantic-density and clip controls. Gray violins show the complete 1,000-sample empirical circular-shift null distributions, dark horizontal segments their 2.5th-97.5th percentiles, short black ticks their means and colored diamonds the observed coefficients. These null intervals are not confidence intervals around the observed coefficients. Displayed q values are BH-FDR across Global plus seven networks. Control and Salience/ventral attention each had q=0.004, whereas the global effect was not significant. d, Clip-specific coefficients for all 13 movie clips. Control was positive in 9/13 clips and survived clip-as-replicate FDR (q=0.026); Salience/ventral attention was positive in 10/13 clips but did not survive this analysis (q=0.096). e, Dynamic-commonality workflow and nuisance controls. WordNet change is an automated feature transition, not a human-annotated narrative boundary. These associations do not establish cortical-layer or causal mechanisms.

The parcelwise separation map derived from stimulus-shared ISFC resembled the separation map from ordinary within-participant MOVIE FC (r = 0.410, p_spin_ = 0.00050; Fig. 4b). Stimulus-locked responses therefore reproduced part of the spatial organization seen during movie viewing. However, ISFC separation was not significantly related to the REST-to-MOVIE ΔSep map (r = 0.128, p_spin_ = 0.127; Fig. S6d), so it did not explain why separation changed between conditions. An earlier post-embedding circular-shift null was not used for inference because its estimated separation increased as alignment with the reference gradient deteriorated (r = −0.36, p = 0.0002; Fig. S6e). This dependence indicates that the null separation was partly driven by unstable gradient orientation rather than by a meaningful null distribution. Descriptively, the final run-balanced analysis showed that ISFC separation varied across networks, with larger values in Somatomotor and Limbic cortex and smaller values in Visual and Default regions (Fig. S6f).

We then tested whether moments of larger semantic change were accompanied by greater depth commonality, that is, whether stimulus-locked activity showed stronger across-participant synchrony in the component shared by the midthickness and white-boundary signals than in the component distinguishing them. WordNet semantic change was shifted by 5 s to approximate the hemodynamic delay and was modeled together with semantic density, MotionEnergy level and change, and clip identity (Fig. 4e). The whole-cortex association was not significant (β = 0.029, q = 0.211). In contrast, semantic change predicted greater depth commonality in Control (β = 0.099, incremental R^2^ = 0.0071, q = 0.0040) and Salience/ventral attention (β = 0.082, incremental R^2^ = 0.0049, q = 0.0040; Fig. 4c). These were the only two outcomes that survived correction across the whole cortex and seven networks (Fig. S7a).

To assess whether these pooled associations generalized across movie content, we re-estimated the controlled semantic-change coefficient separately within each of the 13 clips (Fig. 4d). A positive clip-specific coefficient indicates that, within that clip, moments of larger semantic change were associated with greater depth commonality after controlling for semantic density and MotionEnergy level and change. Control coefficients were positive in 9 clips and remained significant when clips, rather than time points, were treated as the observations (q = 0.026). Salience/ventral-attention coefficients were positive in 10 clips but did not survive that more stringent cross-clip test (q = 0.096). The complete network-by-clip matrix likewise showed substantial variation among movie segments (Fig. S7b). Neither pooled effect was driven by a single clip, because leave-one-clip-out estimates remained positive (Fig. S7c). Predictor correlations did not indicate redundancy between semantic change and the nuisance features (Fig. S7d), and positive coefficients occurred near the prespecified 5-s lag in both networks (Fig. S7e). An example clip illustrates the aligned semantic-change and Control-commonality time series used in the model (Fig. S7f). Control therefore provided the stronger evidence for generalization across movie content; the Salience result was supported by the time-shift null but was less consistent across clips. WordNet change represents an automated lexical-semantic transition, not a human-annotated narrative boundary.

### REST spatial organization generalized across 7 T and 3 T, but magnitude did not

Finally, we asked whether the REST separation pattern established at 7 T could be recovered at 3 T in the 172 participants with complete REST1 LR and RL data. The mean 3 T map (Fig. 5a) and matched 7 T map (Fig. 5b) showed similar cortical topographies. Across 400 parcels, their correlation was r = 0.826 and exceeded the 10,000-spin null (p_spin_ = 0.00030; Fig. 5c). Correspondence remained strong after removing the mean value of each Yeo network from both maps (r = 0.788, p_spin_ = 0.00030; Fig. S10c), indicating that similarity extended beyond the broad ordering of the seven networks.

**Figure 5.**
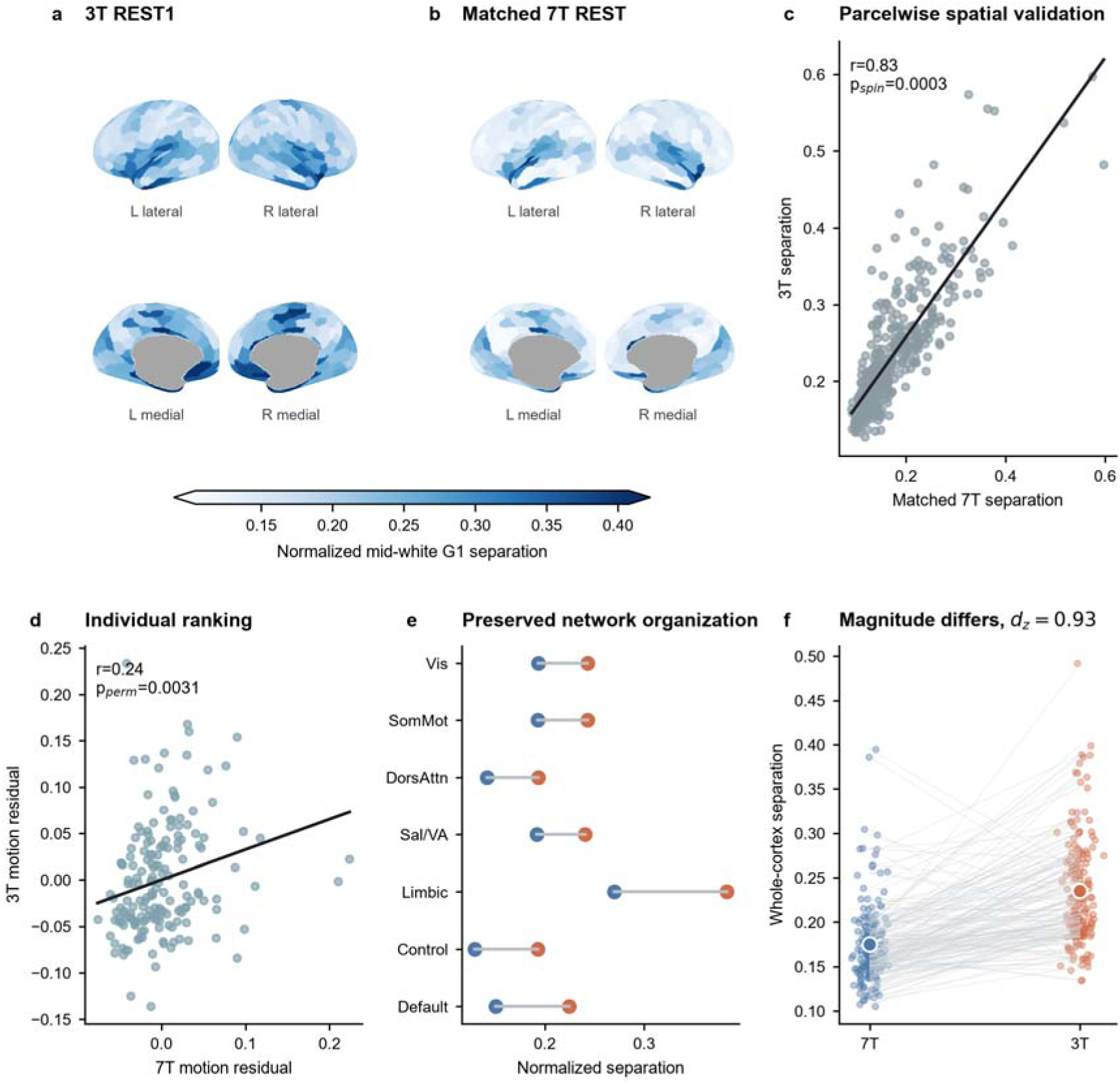
REST depth-gradient organization generalizes from 7T to 3T, but magnitude does not. a,b, Across-participant mean normalized G1 midthickness-to-white-boundary separation maps for 3T REST1 and matched 7T REST in the 172-participant complete-case cohort. Both maps use the same sequential blue scale. To prevent a small number of extreme parcels from compressing the visible cortical pattern, the display range is fixed to the pooled 2nd and 98th percentiles across both maps (0.103-0.407); color-bar extensions denote values outside this range. This clipping affects visualization only and was not applied to any statistical analysis. Gray denotes the medial wall without a Schaefer assignment. c, Spatial correspondence across the same 400 cortical parcels. The black line is the least-squares fit; inference used 10,000 hemisphere-preserving spins to account for cortical spatial autocorrelation (Pearson r=0.826, p_spin_=0.00030). d, Cross-field individual ranking after residualizing each field’s separation estimate for its own motion. Each point is one participant; inference used 10,000 subject-label permutations (r=0.239, p_perm_=0.0031; bootstrap 95% CI 0.106-0.390), indicating modest rather than strong individual generalization. e, Seven-network means. Blue points are 7T, orange points 3T and gray lines join the same network, showing preserved network organization alongside higher 3T values. f, Paired whole-cortex separation for all 172 participants. Gray lines join the same participant, translucent points show individuals and large circles group means. Mean separation was 0.174 at 7T and 0.235 at 3T (mean difference=0.061, paired d_z_=0.935; motion-adjusted p=2.97 × 10^-25^). This is same-participant cross-field validation of REST spatial organization, not independent-cohort replication, numerical equivalence across acquisition conditions or validation of the 7T MOVIE effect.

This spatial result was robust to alternative ways of defining the gradient coordinate system and the number of components. Most participant gradients aligned well to the fixed 7 T reference, although alignment was weaker for the 3 T white-boundary surface and included several low-correlation observations (Fig. S10a). Positive 3 T-7 T map correspondence was observed for fixed-reference participant means, gradients from group-average FC, and a symmetric 3 T-7 T reference, as well as between the two 3 T phase-encoding maps (Fig. S10b). Spatial correlations increased when G2 and G3 were added (G1-G2, 0.88; G1-G3, 0.89), whereas participant-rank correlations remained much smaller (Fig. S10d).

At the individual level, motion-adjusted 3 T and 7 T separation scores were positively but modestly associated (r = 0.239, p_perm_ = 0.0031; bootstrap 95% CI, 0.106-0.390; Fig. 5d). Within 3 T, participant scores from the LR and RL runs correlated at r = 0.56. The two-way absolute-agreement intraclass correlation coefficient for the average of the two runs was 0.71 (ICC(A,2); Fig. S10e). Combining phase-encoding directions therefore improved within-3 T reliability, but the ordering of participants was much less stable across field strengths than the group-average cortical map.

The broad network pattern was preserved: Limbic cortex showed the largest separation at both field strengths, whereas Control and Dorsal-attention regions were lower (Fig. 5e). Nevertheless, every network was shifted upward at 3 T. Whole-cortex separation averaged 0.235 at 3 T and 0.174 at 7 T, a paired difference of 0.061 (d_z_ = 0.935; motion-adjusted p = 2.97 × 10^-25^; Fig. 5f). Motion showed limited positive associations with separation and with the field-strength difference, but adjustment did not remove the field effect (Fig. S10f). The 3 T analysis therefore supports generalization of the REST spatial pattern in the same participants, but not numerical equivalence, a stable cross-field individual trait, or validation of the 7 T MOVIE result.

## DISCUSSION

This study shows that cortical sampling position is not a neutral implementation detail, but a measurable dimension of macroscale functional-connectivity organization. Signals sampled at midthickness and the gray-white boundary were strongly related, yet their connectomes occupied nonredundant positions in gradient space. The separation between those positions was spatially structured at rest, changed selectively during movie viewing, and was preserved topographically across 7 T and 3 T REST acquisitions. At the same time, effect magnitude and participant ranking were substantially less stable than cortical topology. These findings therefore distinguish three properties that are often conflated: where a cross-surface difference is expressed across cortex, how large that difference is under a given acquisition, and how reliably individuals can be ordered by it. The principal result is not that the two surfaces isolate separate biological compartments, but that a small shift in where BOLD time series are sampled reveals a reproducible, state-sensitive organization of whole-connectome geometry.

The result extends evidence that the gray-white boundary has a distinctive functional position. Our previous work showed that activity and connectivity can change sharply across this boundary and that the contrast is spatially organized rather than reducible to a uniform gray-versus-white offset (Li et al., 2025). Here, that local boundary contrast is linked to a global property: the relative placement of two sampling surfaces within a low-dimensional connectome manifold. This link matters because surface placement and cortical geometry can systematically alter which tissue and signals contribute to a measurement, particularly in thin or highly folded cortex (Coalson et al., 2018). Yet the present effect cannot be summarized as a fixed geometric bias. A fixed offset should be similar across brain states, whereas movie viewing changed cross-surface separation in a network-and parcel-specific manner. The boundary is therefore best treated as a functionally distinctive measurement location whose relationship to a more superficial cortical sample depends on the ongoing organization of connectivity, without implying that either sample is anatomically pure or layer specific.

Movie viewing reorganized this relationship on top of a stable macroscale scaffold. The broad cortical arrangement of separation was retained from REST to MOVIE, but the global reduction in separation was not uniform: Visual, Limbic and Default regions converged most consistently, whereas Salience/ventral-attention cortex showed a G1-specific tendency toward differentiation. This combination of stability and flexibility accords with evidence that task states reconfigure intrinsic network organization without replacing it (Cole et al., 2014; Gratton et al., 2018). It also complements work showing that the principal cortical gradient flexes with task demands (Gao et al., 2022) and that gradients measured during naturalistic processing retain hierarchical structure while varying by modality and content (Samara et al., 2023). Our contribution is to show that such reconfiguration also has a sampling-position dimension: state effects alter not only the location of cortical parcels along a gradient, but also the distance between connectome representations obtained from nearby cortical surfaces. The network pattern further argues against interpreting the global MOVIE convergence as simple signal homogenization. If that were the sole explanation, comparable changes would be expected across systems and gradient dimensions; instead, the direction and dimensional specificity depended on network affiliation.

The decomposition analyses clarify what changed mathematically while setting an important limit on mechanistic interpretation. Exact Shapley attribution assigned the REST-to-MOVIE change to coordinated movement of both surfaces rather than to a single dominant surface (Shapley, 1953). Direct counterfactuals reached the same conclusion: neither holding midthickness fixed nor holding the white-boundary surface fixed reproduced the observed state-change map. Shapley values here are algebraic allocations of a two-input distance change, not evidence that either surface drives the other. The result is instead consistent with a joint reconfiguration of two related connectome representations. Analyses in the original functional-connectivity space provided a convergent measurement check: changes in cross-surface FC-profile similarity tracked changes in gradient separation, reducing the likelihood that the finding was created solely by nonlinear embedding. Because gradients compress connectivity into dominant axes (Margulies et al., 2016; Vos de Wael et al., 2020), this agreement across gradient and edge-profile descriptions supports the robustness of the phenomenon while preserving the distinction between corroboration and independent replication.

Naturalistic stimulation provides a particularly informative setting for observing this state dependence. Unlike short event-related or block designs interleaved with frequent rest, the HCP movie runs contain extended, uninterrupted stimulation; correlations can therefore be estimated within a sustained task context with less contamination from repeated task-rest transitions. Natural movies also synchronize cortical responses across viewers (Hasson et al., 2004) and support shared, time-varying network configurations during narrative comprehension (Simony et al., 2016). Consistent with this literature, both sampling surfaces showed strong stimulus-locked ISFC, and ordinary MOVIE separation corresponded to the separation derived from stimulus-shared connectivity. ISFC nevertheless did not account for the MOVIE-minus-REST separation map, so shared timing and state-dependent cross-surface reconfiguration should be regarded as complementary rather than interchangeable phenomena (Nastase et al., 2019). The semantic analysis further localized a modest content-related component. WordNet-defined semantic change predicted a greater common-mode contribution most robustly in Control cortex, whereas the Salience/ventral-attention effect was less stable across clips. This pattern is compatible with the role of control systems in organizing context-dependent cognition and with distributed semantic representations during natural viewing and speech (Badre et al., 2005; Huth et al., 2016, 2012). Importantly, WordNet change measures lexical-semantic displacement, not narrative event boundaries; the result therefore identifies one restricted correlate of cross-surface synchrony rather than a complete account of movie structure.

The venous and myelin-sensitive analyses constrain two straightforward anatomical explanations. Separation and its state change did not preferentially concentrate in parcels with greater VENAT venous probability, and the principal findings persisted after excluding high-probability parcels. Because large veins and intracortical venous drainage can influence the spatial specificity of gradient-echo BOLD measurements (Huck et al., 2019; Kay et al., 2019; Markuerkiaga et al., 2016), these null results weigh against a simple group-topographic account in which mapped venous density alone produces the cortical pattern. Likewise, weak or absent associations with T1w/T2w-derived myelin measures argue against explaining the separation map as a direct reflection of one conventional myelin-sensitive contrast (Glasser and van Essen, 2011; Hagiwara et al., 2018). These tests do not exclude vascular or microstructural contributions to either sampled signal. Rather, they show that the reported topology and state change are not readily predicted by the available group venous atlas or parcelwise T1w/T2w measure.

The matched 3 T-7 T analysis further separates robust spatial organization from acquisition-dependent scale. REST separation maps were strongly correlated across field strengths, remained similar after removing network means, and preserved the same broad network ordering. By contrast, absolute separation was larger at 3 T and cross-field participant correspondence was modest. Field strength, spatial resolution and sequence properties jointly alter BOLD sensitivity, noise and partial-volume mixing (Triantafyllou et al., 2005; Uğurbil et al., 2013; Vu et al., 2017). The present data therefore support cross-acquisition generalization of group-level REST topology, not numerical equivalence of effect size. They also do not constitute an independent-cohort replication or a validation of the MOVIE effect at 3 T. More generally, the contrast between strong map correspondence and limited individual correspondence reinforces the need to report spatial reproducibility, magnitude agreement and rank-order reliability as separate validation targets.

Three limitations define the appropriate scope of these conclusions. First, spatial resolution remains insufficient to treat midthickness and white-boundary time series as isolated tissue or laminar measurements. They are nearby spatial samples from a partially overlapping hemodynamic field, and vascular point-spread and partial-volume effects limit anatomical specificity. Second, individual reliability is limited. Group maps and network patterns were robust, but cross-run and especially cross-field participant rankings were only moderate, consistent with the broader observation that stable individual functional-connectivity estimates require substantial data and careful reliability assessment (Birn et al., 2013; Gordon et al., 2017; Noble et al., 2019). The present measure is consequently better supported as a group-level organizational feature than as an individual biomarker. Third, the semantic feature space was limited. WordNet provides a principled lexical taxonomy (Miller, 1995), but it does not capture the full range of perceptual, social, affective and narrative structure represented during naturalistic viewing, nor the multiscale event organization of continuous experience (Baldassano et al., 2017; Huth et al., 2012). Broader, preregistered feature sets will be required to determine which aspects of movie content modulate cross-surface commonality.

In conclusion, functional-connectivity gradients depend not only on brain state and analytical scale, but also on where the underlying BOLD signal is sampled relative to the cortical boundary. By separating state-dependent changes in spatial organization from acquisition-dependent changes in magnitude, these findings establish cortical sampling position as an interpretable measurement dimension of macroscale functional organization.

## Data Availability

The HCP-Y data used in this study are available in the HCP database https://www.humanconnectome.org/

## Code Availability

Custom MATLAB and Python scripts developed for this study to perform the analyses and generate the figures are available from the corresponding author upon reasonable request. Analyses were implemented in MATLAB R2026a and Python 3.14 using Connectome Workbench v1.5.0 for volume-to-surface sampling, and the MATLAB implementation of BrainSpace for functional-gradient estimation and alignment. Python-based statistical analysis, neuroimaging I/O, surface visualization, and figure generation used NumPy, SciPy, pandas, NiBabel, Nilearn, and Matplotlib.

## ACKNOWLEDGMENTS

This work was supported by the National Institutes of Health (NIH) grants R01 NS113832 (JG) and R01 NS129855 (ZD).

Data of young adults were provided by the Human Connectome Project, WU-Minn Consortium (Principal Investigators: David Van Essen and Kamil Ugurbil; 1U54MH091657), funded by the 16 NIH Institutes and Centers that support the NIH Blueprint for Neuroscience Research; and by the McDonnell Center for Systems Neuroscience at Washington University.

## AUTHOR CONTRIBUTIONS

M.L.: Writing, Visualization, Validation, Software, Methodology, Investigation, Conceptualization. Z.D.: Supervision and Funding acquisition. J.C.G: Supervision and Funding acquisition.

