## Supplemental Figures for "Functional connectivity gradients depend on cortical sampling position and brain state"

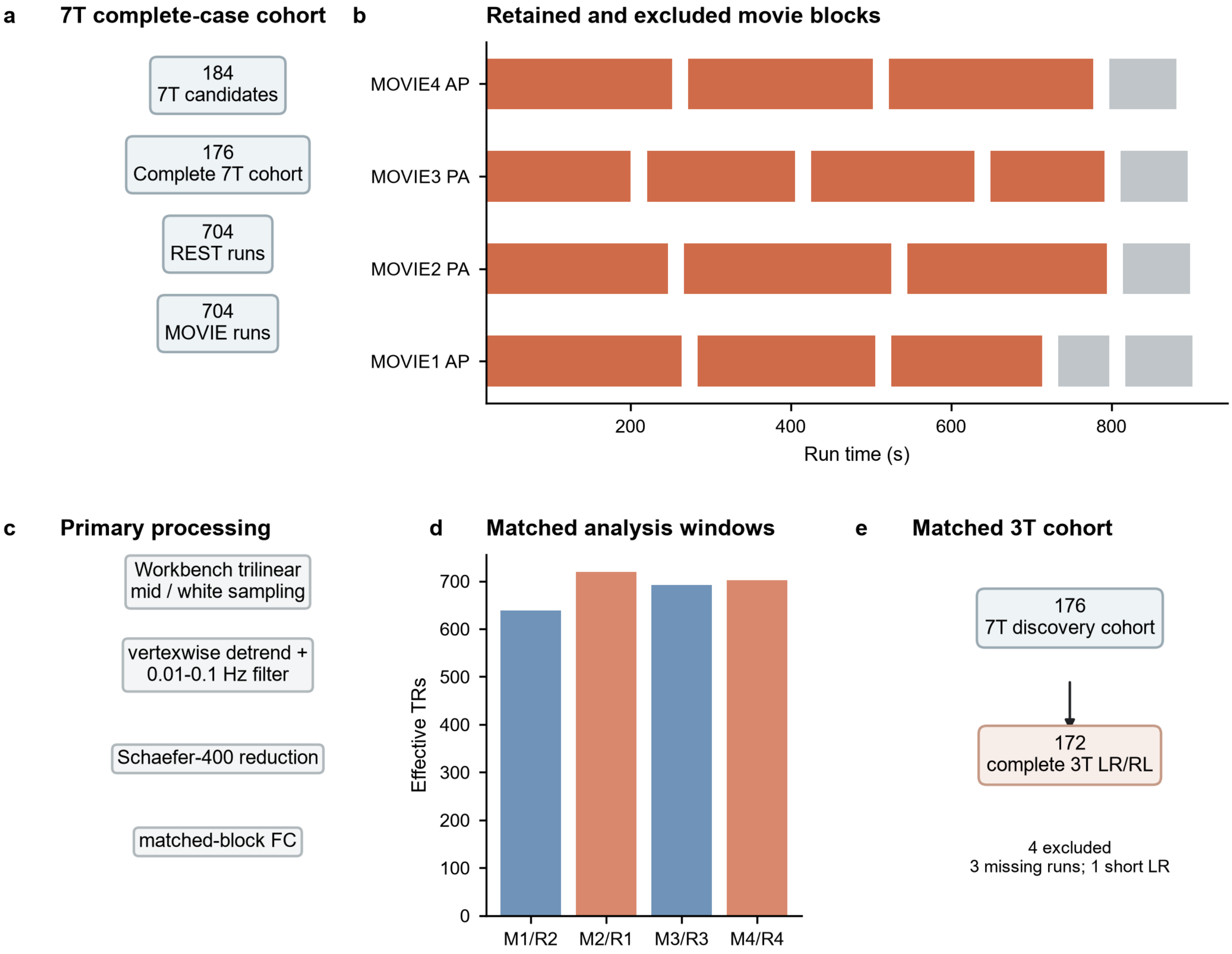


**Figure S1.** Cohort, acquisition and matched-block design. a, Complete-case derivation of the 7T discovery cohort. Of 184 candidates, 176 participants had all four REST and four MOVIE runs, yielding 704 runs per state. b, Official timing of the four movie runs. Coral blocks were retained in the primary state comparison and gray blocks excluded; the excluded material includes the 64-s clip and repeated Vimeo content. AP and PA denote phase-encoding direction. c, Primary processing sequence: subject-specific Workbench trilinear sampling at midthickness and the gray-white boundary, vertexwise detrending and 0.01-0.1-Hz filtering, Schaefer-400 parcel reduction and duration-matched functional connectivity. d, Effective samples in the four prespecified MOVIE-REST pairings after applying the common [block start+5 s, block end) window. M and R denote MOVIE and REST runs; bars show retained TRs. Valid blocks were duration-weighted within run and runs received equal weight. e, Derivation of the matched 3T validation cohort. Of the 176 7T participants, 172 had complete usable 3T REST1 LR and RL data; three lacked a required run and one LR run was too short. Stimulus-shared connectivity uses separate effective-duration clip weights shown in Extended Data Figure 6. Source data are provided with the figure.

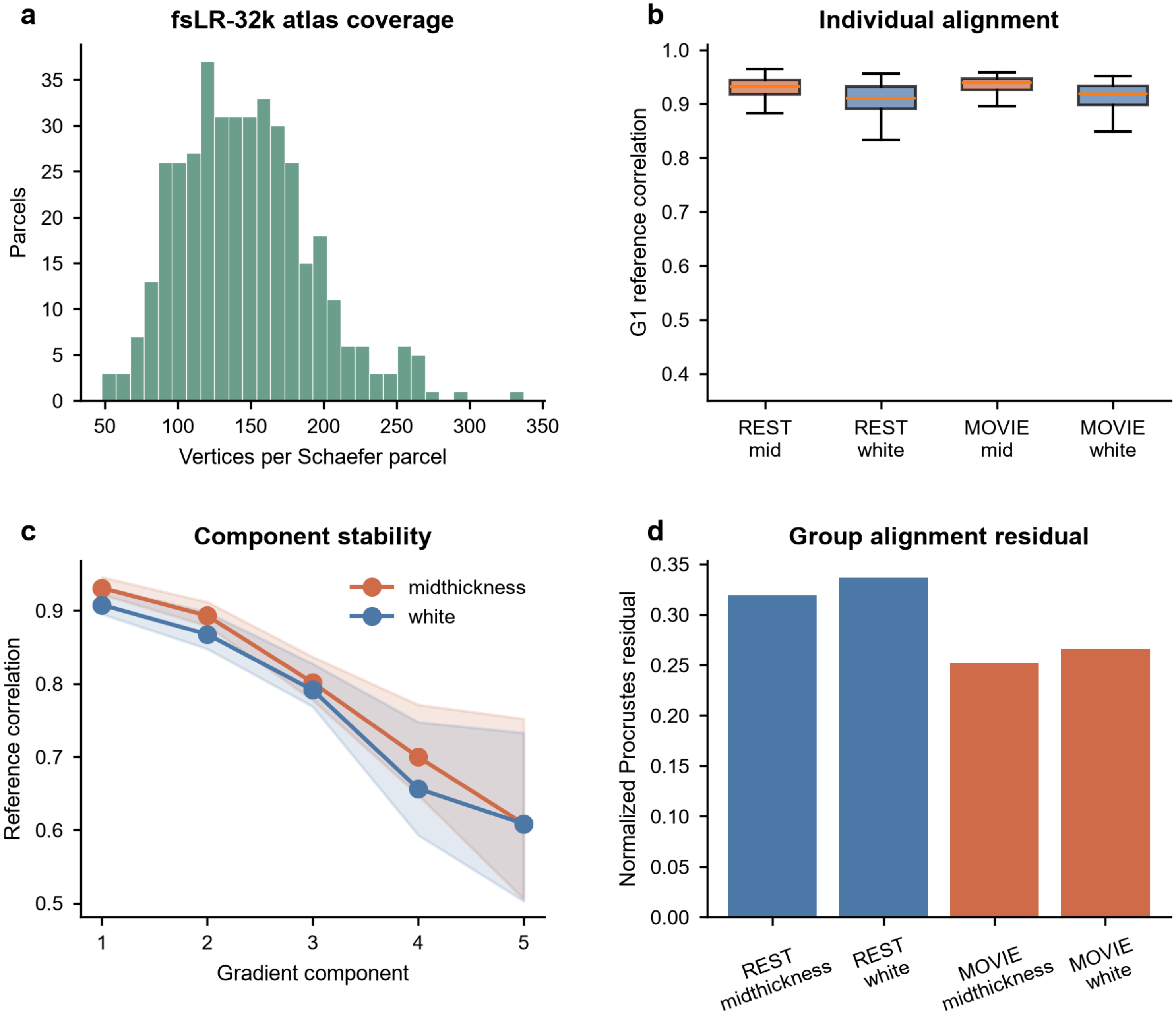


**Figure S2.** Surface sampling and gradient-alignment quality control. a, Number of valid fsLR-32k vertices contributing to each Schaefer-400 cortical parcel after subject-specific midthickness and gray-white-boundary sampling; distributions summarize cortical coverage rather than a functional effect. b, Participant-level Pearson correlation between each aligned G1 and the locked common G1 reference, shown separately by state and sampled surface for the 176-participant 7T cohort. c, Reference-correlation distributions for G1-G5. Higher values indicate that the estimated component preserves the orientation of the common reference; the comparison establishes strong G1-G2 alignment, broadly usable G3 alignment and weaker higher-component stability. d, Group-level Procrustes residuals after alignment; smaller values indicate closer agreement with the reference embedding. The primary separation analysis uses scale-normalized G1. G1-G2 and G1-G3 distances are sensitivity analyses, whereas G4-G5 are not used for biological interpretation. These panels assess sampling and numerical alignment quality and do not constitute evidence for a state effect. Source data are provided with the figure.

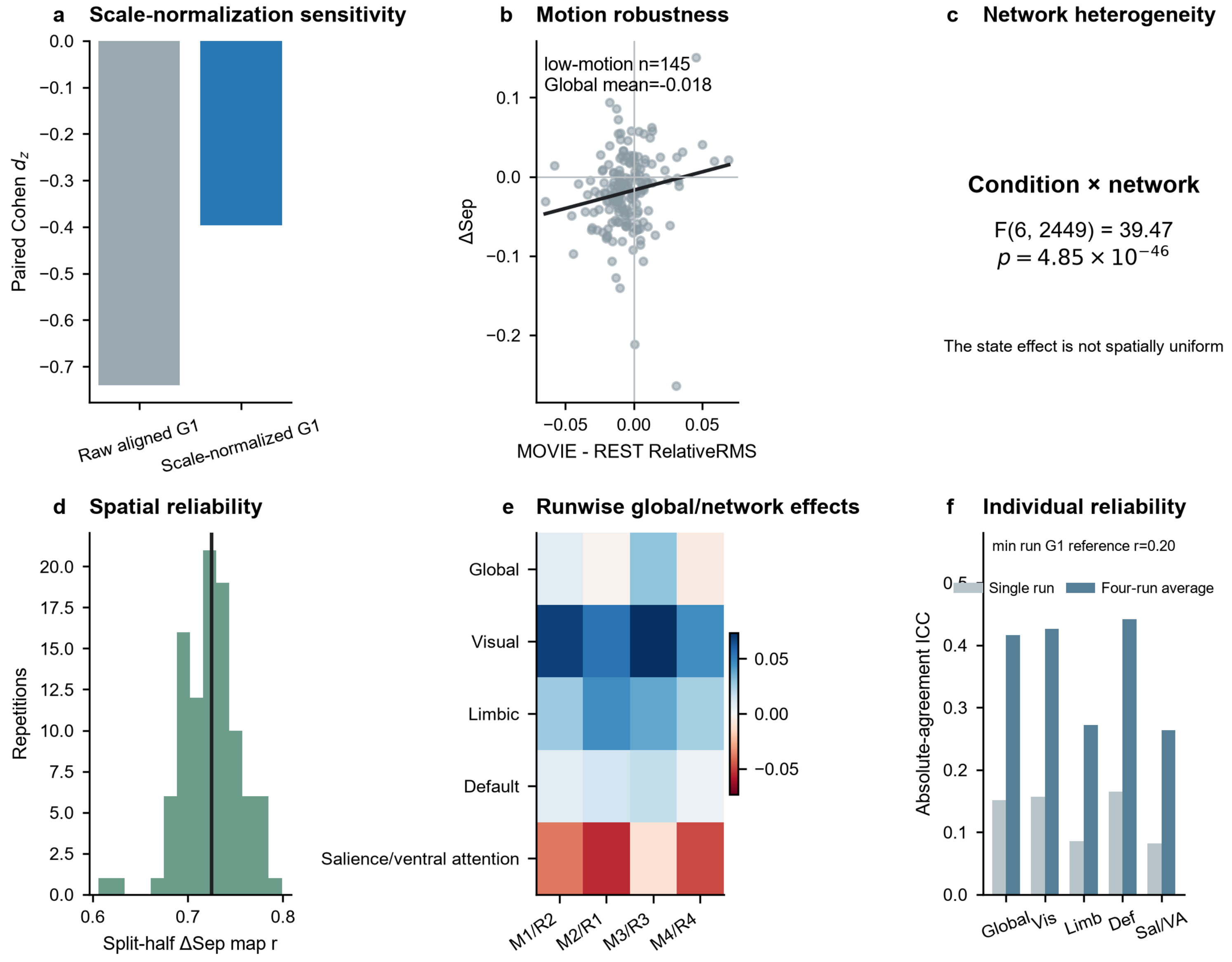


**Figure S3.** Robustness, scale contribution and runwise reliability of the primary state effect. a, Paired Cohen $d_z$ for the raw aligned-G1 distance and the within-subject scale-normalized G1 distance in 176 participants. Both estimates indicate MOVIE-related convergence (raw $d_z$=-0.74; normalized $d_z$=-0.40), showing that scale contributes to but does not explain the effect. b, Participant ΔSep versus MOVIE-minus-REST RelativeRMS. Points are participants and the line is the least-squares fit; inset values report the prespecified low-motion subset for which mean RelativeRMS in both states was below 0.15. c, Omnibus condition-by-network interaction from the repeated-measures mixed model. The displayed F statistic and degrees of freedom test whether the state effect is spatially uniform across seven networks. d, Distribution of independently rebuilt split-half correlations for the parcelwise ΔSep map; the vertical line is the median. e, IntegrationGain, defined as REST minus MOVIE separation, across four duration-matched run pairs. Blue and red indicate positive convergence and negative differentiation, respectively. The global effect varies across pairs, whereas the focal network directions are more consistent; pairs reuse the same participants and are not independent replications. f, Absolute-agreement ICC for single-run scores and four-run averages. The annotation reports the minimum runwise G1 reference correlation as an alignment boundary. Reliability improves with averaging but remains insufficient for a stable individual-trait claim. Source data are provided with the figure.

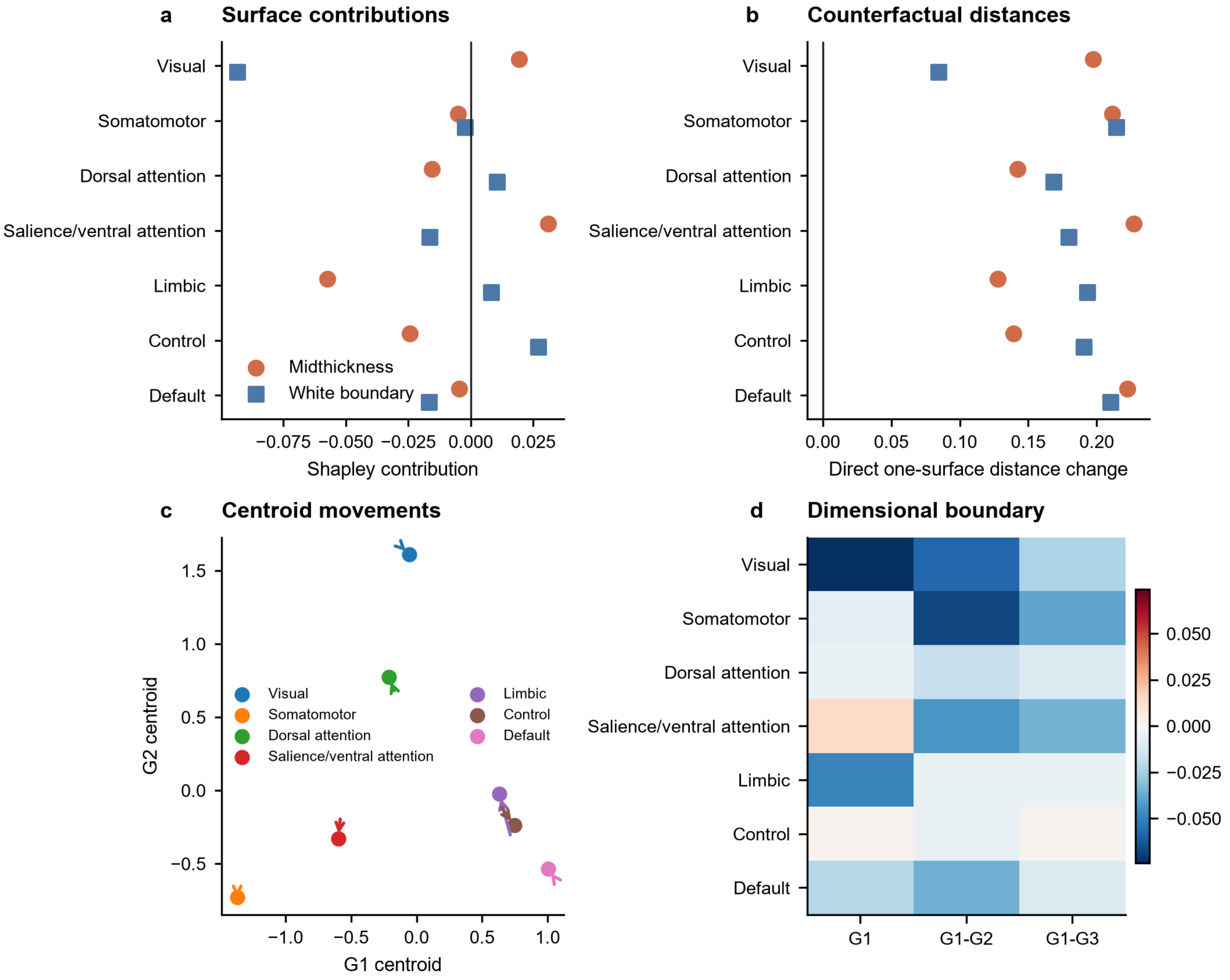


**Figure S4.** Complete surface-contribution and dimensionality analyses. a, Midthickness Shapley contributions (orange circles) and white-boundary contributions (blue squares) to G1 ΔSep for all seven networks. Negative values reduce G1 separation and promote convergence; positive values increase separation and promote differentiation. The two contributions sum exactly to ΔSep. b, Direct one-surface-at-a-time counterfactual changes. Orange circles show $D_{10}$-$D_{00}$, obtained by changing midthickness while holding the white boundary at REST; blue squares show $D_{01}$-$D_{00}$, obtained by changing the white boundary while holding midthickness at REST. Positive values mean that changing either surface alone increases its distance from the other surface's REST representation. c, REST-to-MOVIE movement of each network's midthickness centroid in aligned G1-G2 space; arrows indicate state direction and colors Yeo-7 network identity. Complete midthickness and white-boundary REST/MOVIE coordinates are provided as Source Data. d, Mean MOVIE-minus-REST distance change using G1 alone, Euclidean G1-G2 distance and Euclidean G1-G3 distance. Blue denotes convergence and red differentiation. Salience/ventral-attention differentiation is specific to G1 and should not be described as whole-manifold differentiation. These are movements in functional-gradient space, not anatomical movement or causal direction between cortical depths.

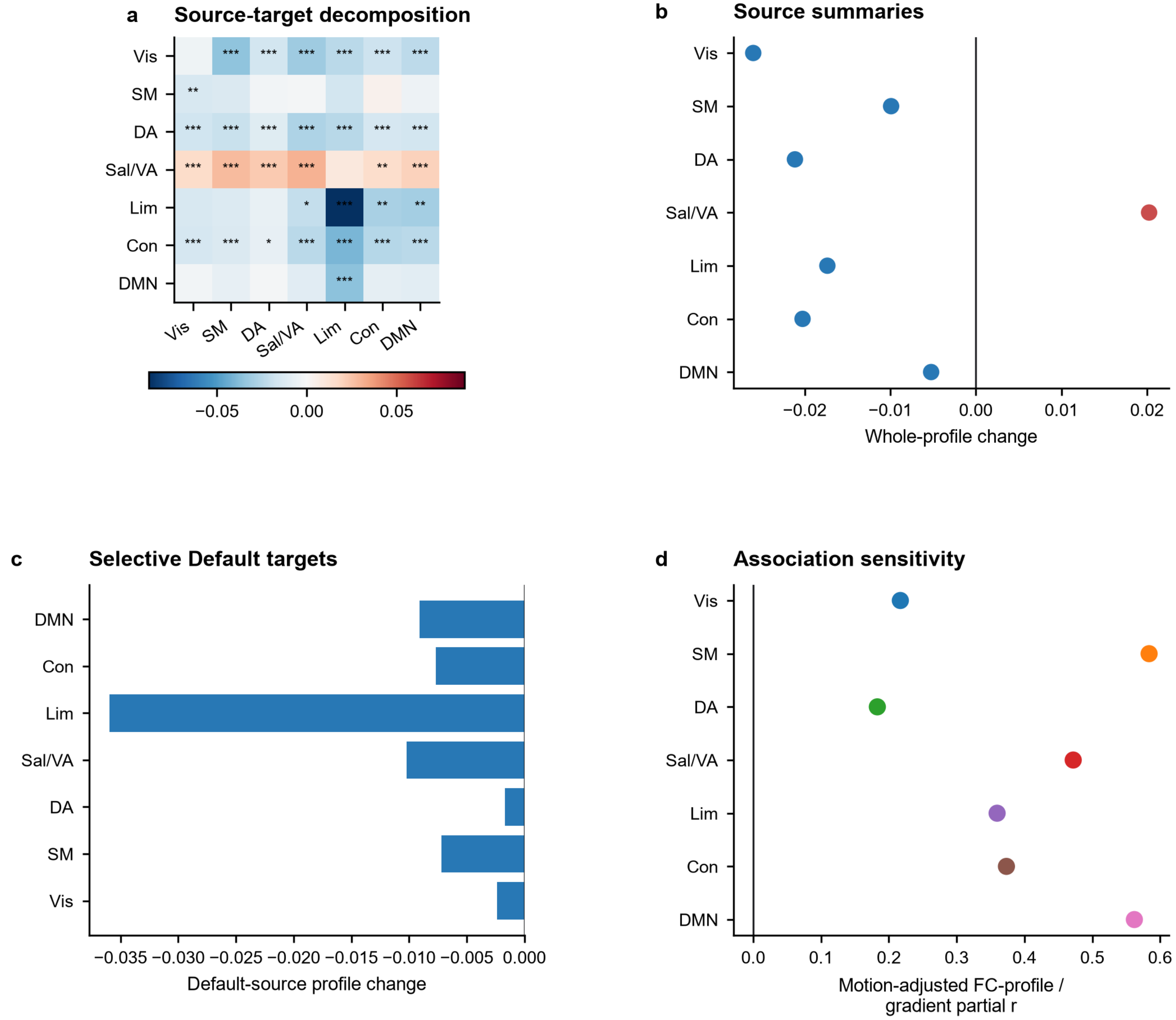


**Figure S5.** Full FC-profile decomposition and association sensitivity. Network abbreviations are Vis, visual; SM, somatomotor; DA, dorsal attention; Sal/VA, salience/ventral attention; Lim, limbic; Con, control; and DMN, Default. a, Complete seven-source-by-seven-target matrix of MOVIE-minus-REST cross-depth FC-profile distance. Rows identify source networks and columns target networks. Blue values indicate that standardized midthickness and white-boundary FC profiles become more similar during MOVIE; red values indicate greater dissimilarity. Asterisks mark motion-adjusted BH-FDR across all 49 source-target tests (*q<0.05, **q<0.01, ***q<0.001). b, Whole-profile MOVIE-minus-REST change for all seven source networks. These values are calculated directly from each source parcel's complete 399-target standardized FC profile and are not averages of the seven cells in panel a. c, Target-resolved effects for Default-network sources. The Default-to-Limbic profile converges despite the null Default whole-profile summary, demonstrating a selective rather than global Default reconfiguration. Bar color encodes convergence or differentiation and the vertical line no change. d, Participant-level association between whole-profile change and G1 separation change after residualizing both variables for MOVIE-minus-REST motion difference. Points are network partial Pearson correlations; positive values mean that stronger profile convergence accompanies stronger gradient convergence, and stronger differentiation accompanies stronger gradient differentiation. The correspondence is embedding-independent support for the gradient pattern but does not establish causality. Complete participant-level source-target data are provided with the figure.

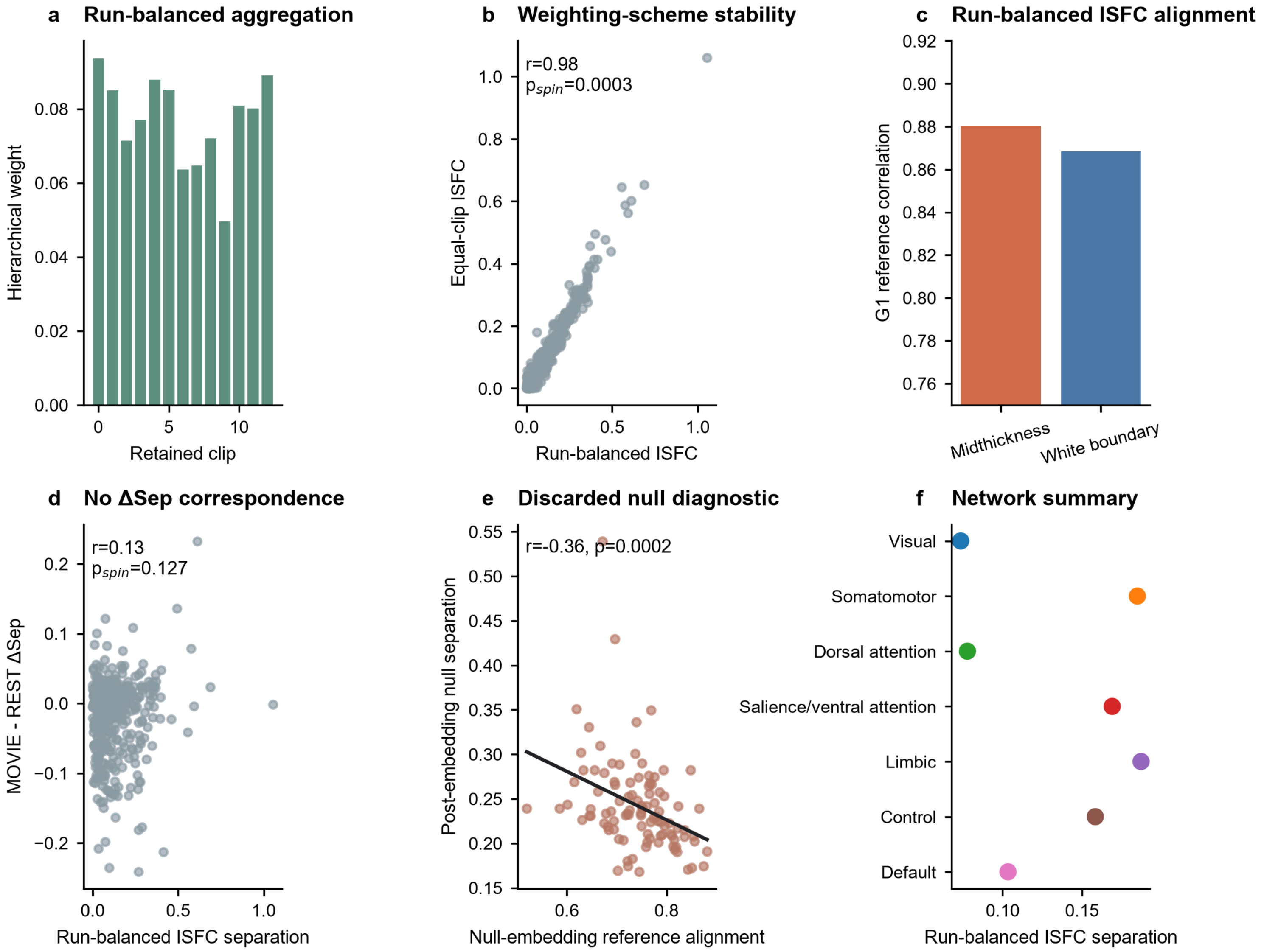


**Figure S6.** Run-balanced ISFC validation and diagnostic comparisons. All inferential results use the final run-balanced analysis. The earlier equal-clip aggregation is shown only to test whether the spatial map is robust to the weighting scheme, and the earlier post-embedding circular-shift null is shown only to document why that null procedure was discarded. a, Hierarchical aggregation weights for retained movie clips. Within each run, clips are weighted by effective duration after the 5-s onset buffer; the four runs then receive equal total weight. b, Parcelwise comparison of run-balanced ISFC separation with the alternative equal-clip aggregation. Each point is one of 400 cortical parcels; the map remains highly similar (Pearson r=0.979, $p_{spin}$=0.00030), indicating that weighting changes magnitude more than topology. c, Alignment quality of the run-balanced midthickness and white-boundary G1 embeddings relative to the locked common reference (r=0.880 and 0.868). d, Run-balanced ISFC separation versus ordinary MOVIE-minus-REST ΔSep. The association is not significant (r=0.128, $p_{spin}$=0.127), so stimulus-shared ISFC topology should not be described as explaining the state-change map. e, Post-embedding circular-shift null separation versus the null embedding's reference alignment. Their dependence shows that this discarded null statistic is confounded by unstable gradient orientation and is not used for inference. f, Run-balanced ISFC separation summarized across the seven Yeo networks; points are network means. The significant ISFC-to-ordinary-MOVIE topology is shown in Main Figure 4 and is not duplicated here. Source data are provided with the figure.

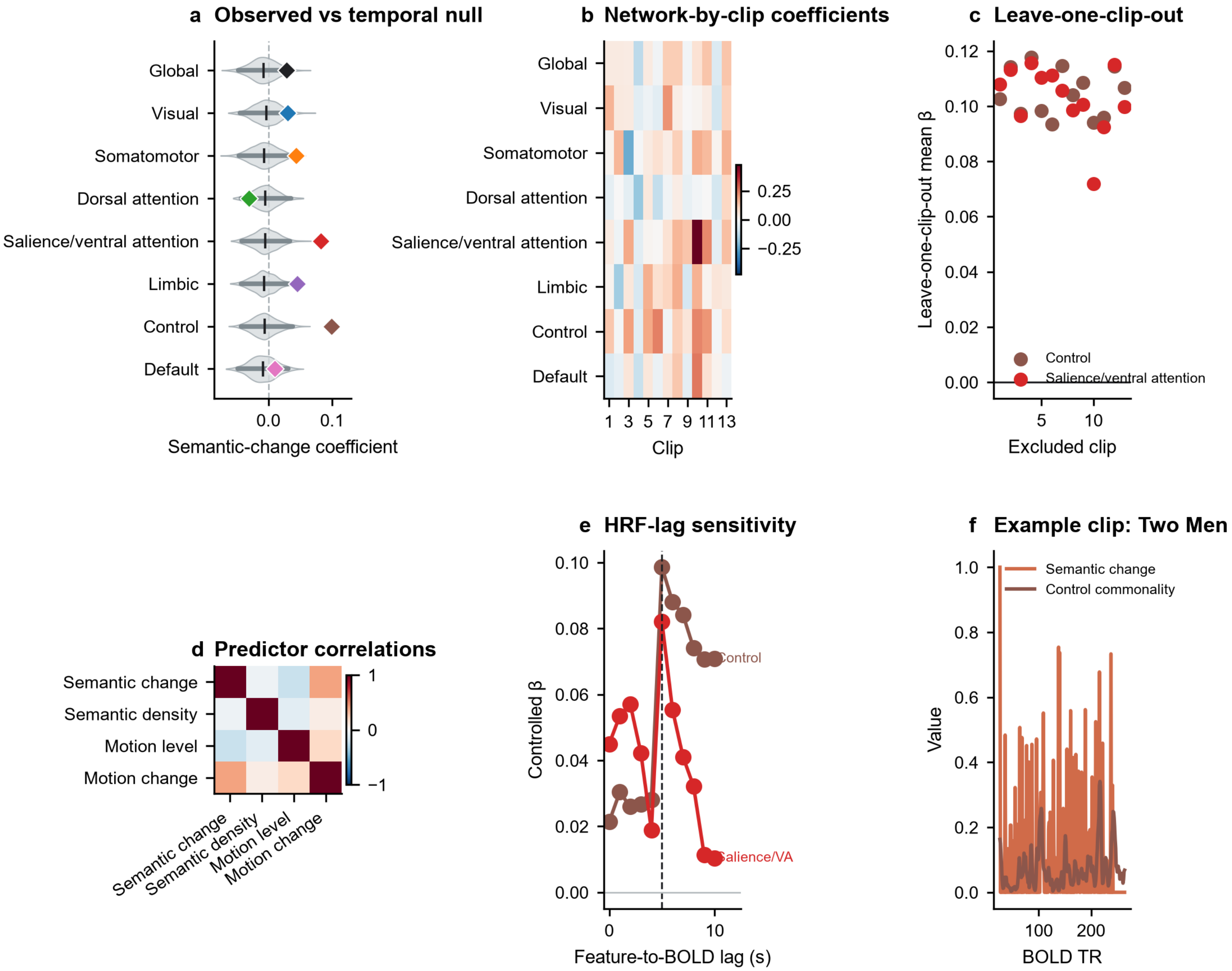


**Figure S7.** Robustness of the semantic-depth commonality model. a, Semantic-change coefficients for the global cortex and seven Yeo networks. Consecutive-TR WordNet change is shifted by 5 s and entered with MotionEnergy level, MotionEnergy change, semantic density and clip fixed effects. Positive coefficients indicate greater common-mode relative to differential-mode synchrony after larger semantic transitions. Gray violins show the complete 1,000-sample empirical circular-shift null distributions, dark horizontal segments their 2.5th-97.5th percentiles, short black ticks their means and colored diamonds the observed coefficients. These null intervals are not confidence intervals around the observed coefficients. FDR covers Global plus seven networks. b, Coefficients for every network and each of 13 movie clips. Blue denotes negative and red positive effects; clip estimates are heterogeneity checks, not independent participant replications. c, Mean Control and Salience/ventral-attention coefficient after excluding each clip in turn. A consistently positive series indicates that no single clip determines the pooled direction. d, Pearson correlations among clip-standardized semantic change, semantic density, MotionEnergy level and MotionEnergy change. The fixed -1 to 1 scale reveals predictor dependence but is not an outcome test. e, Controlled coefficients across feature-to-BOLD lags; the dashed line marks the prespecified 5-s shift. Lag curves are sensitivity analyses rather than multiple independent confirmatory tests. f, Example Two Men time series showing aligned semantic change and Control depth commonality at BOLD TR resolution. Control provides the strongest clip-as-replicate evidence; Salience is significant under time-preserving permutation inference but weaker across clips. WordNet change is not a human-annotated narrative boundary. Source data are provided with the figure.

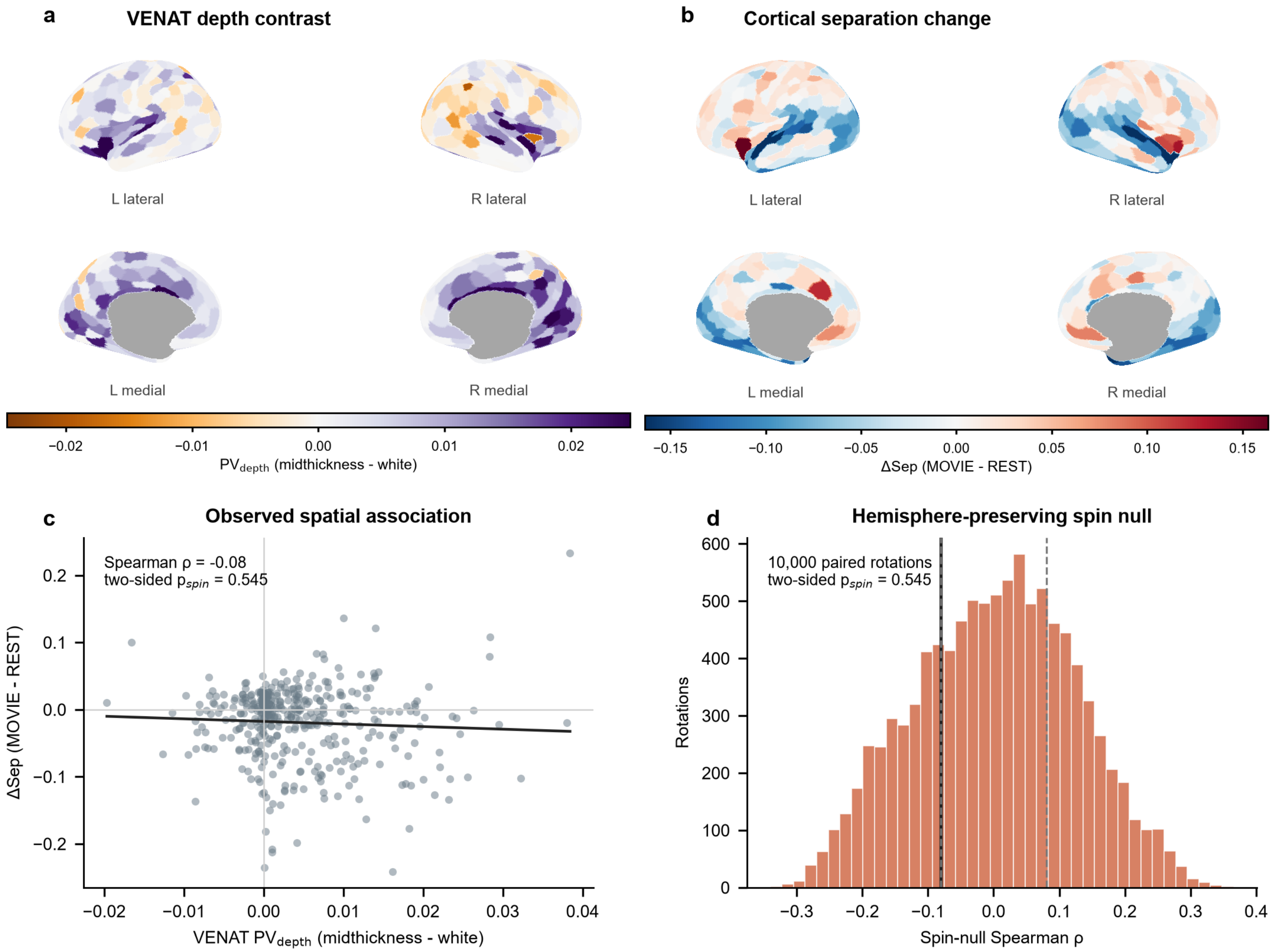


**Figure S8.** VENAT vascular sensitivity analysis. a, Group VENAT atlas estimate of midthickness-minus-white-boundary visible-vein partial-volume contrast, reduced to the Schaefer-400 parcel grid. Positive values indicate greater estimated venous partial volume at midthickness than at the white boundary. b, Mean parcelwise ΔSep, defined as normalized MOVIE separation minus REST separation, on the identical parcel grid. Negative values indicate cross-depth convergence and positive values differentiation. c, Parcelwise association between VENAT depth contrast and ΔSep; each point is one cortical parcel and the line summarizes the monotonic relationship. d, Null distribution from 10,000 hemisphere-preserving spins. The vertical marker is the observed Spearman correlation (ρ=-0.080); its position within the null gives $p_{spin}$=0.545. The null result argues against a simple explanation in which the group-average spatial distribution of visible-vein depth contrast determines the ΔSep map. It does not eliminate vascular contributions, subject-specific vascular anatomy or effects not represented by the group atlas. Source data and the complete spin null are provided with the figure.

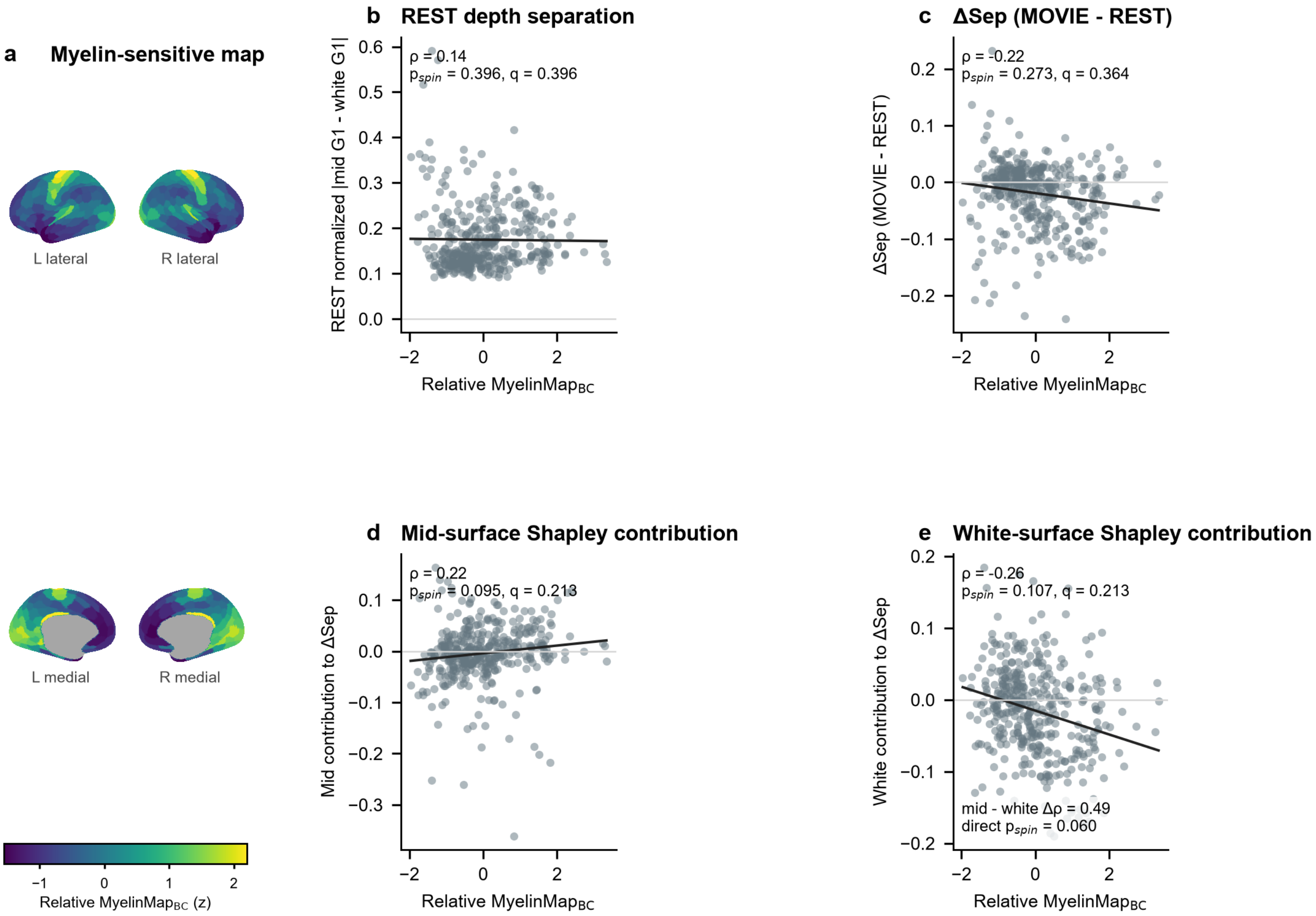


**Figure S9.** Myelin-sensitive structural context of cross-depth gradient organization. a, Group relative MyelinMap$_{BC}$, a T1w/T2w-derived myelin-sensitive contrast rather than a direct measure of cortical myelin content. b-e, Parcelwise spatial associations between this structural map and REST normalized G1 separation, MOVIE-minus-REST ΔSep, the midthickness Shapley contribution and the white-boundary Shapley contribution. Each point represents a cortical parcel; fitted lines are descriptive. Negative ΔSep denotes convergence, whereas the signs of $\varphi_{mid}$ and $\varphi_{white}$ denote each sampled surface's contribution to convergence or differentiation. Primary inference used 10,000 hemisphere-preserving spins to preserve cortical spatial autocorrelation and BH-FDR across the four prespecified spatial tests. None of the primary associations survived correction. The paired-spin comparison of the two surface-contribution directions was suggestive but not significant (p=0.060) and is reported as a trend only. Secondary quadratic and network models, together with subject-specific versus leave-one-out topology analyses, are sensitivity results and do not support an individual-specific myelin mechanism. Source data, spin summaries and secondary-model outputs are provided with the figure.

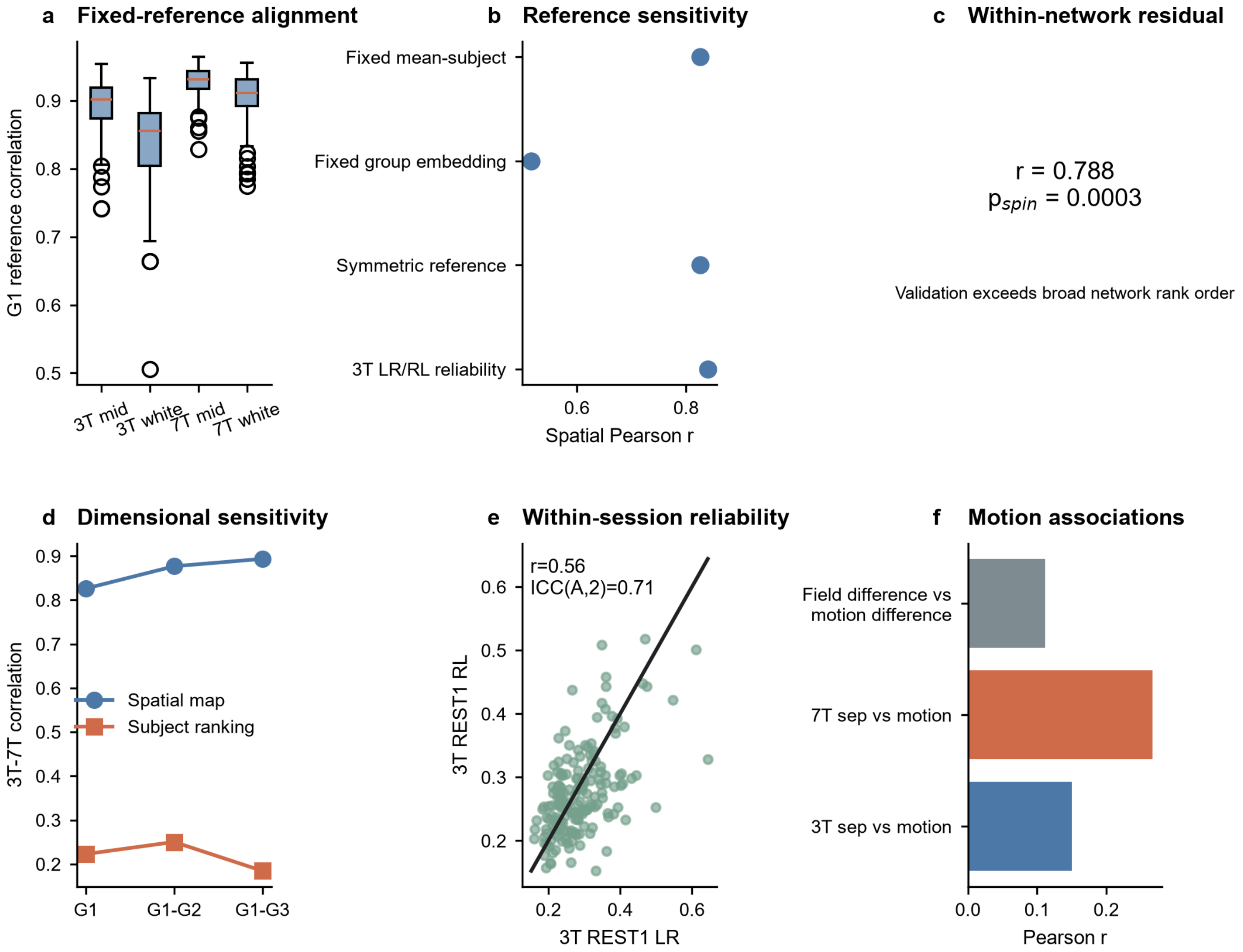


**Figure S10.** Quality control and sensitivity of the 3T-to-7T REST validation. a, Participant-level correlation of aligned G1 with the fixed 7T common reference for 3T REST1 and matched 7T REST, shown separately for midthickness and white-boundary embeddings in the 172-participant cohort. Boxes summarize distributions and center lines medians; higher values indicate stronger reference alignment. b, Spatial Pearson correlations for the primary fixed-reference mean-subject maps, a fixed-reference group-embedding sensitivity, a symmetric-reference sensitivity and 3T REST1 LR/RL reliability. Points summarize 400-parcel map correspondence; spatial inference uses hemisphere-preserving spins. c, Primary 3T-to-7T correspondence after removing Yeo-7 network means from both maps (r=0.788, $p_{spin}$=0.00030), showing that validation extends beyond broad network rank order. d, Cross-field correlation using G1, Euclidean G1-G2 and Euclidean G1-G3 separation. Circles denote group-map spatial correspondence and squares motion-residualized participant ranking; strong spatial validation coexists with modest individual generalization. e, 3T REST1 LR versus RL participant estimates. The black line is identity; Pearson r=0.56 and absolute-agreement ICC(A,2)=0.71 quantify within-session reliability of the two-direction average. f, Pearson associations of 3T separation, 7T separation and their field difference with the corresponding motion measures; the vertical line denotes zero. These are same-participant REST QC and sensitivity analyses, not an independent-cohort replication or validation of the MOVIE effect. Source data are provided with the figure.
